# Association between the skin microbiome and MHC class II diversity in an amphibian

**DOI:** 10.1101/2023.04.12.536591

**Authors:** M Cortazar-Chinarro, A Richter-Boix, P Rodin-Mörch, P Halvarsson, JB Logue, A Laurila, J Höglund

## Abstract

It has become clear that the microbiome plays an important role in determining host health, diseases, and phenotypic variation. There is increasing evidence that the microbiome influences host fitness and its adaptation to the environment is changing our thinking on host-microbe interactions. However, it remains unclear how a host genotype shapes its microbiome. Here, we explored how genetic background and evolutionary history influence associated microbiome in amphibian populations. We studied how skin bacterial diversity is associated with the Major Histocompatibility Complex (MHC) class II exon 2 diversity in 12 moor frog populations belonging to two geographical clusters that show signatures of past and ongoing differential selection patterns. We found that bacterial alpha-diversity remained similar between the two clusters, while MHC haplotype-supertypes and genetic diversity differed between the clusters. Bacterial alpha-diversity was positively correlated with expected MHC heterozygosity and negatively with MHC nucleotide diversity. We also found that bacterial community composition differed significantly between the two geographic clusters and between specific MHC supertypes. These findings further suggest that population historical demographic events influence hologenomic variation and provide new insights into how immunogenetic host variability and microbial diversity may jointly influence host fitness with consequences for disease susceptibility and population persistence.

## Introduction

All complex organisms carry numerous microbes forming diverse microbial communities in many organs including the skin, lungs, and gut (Antwis, Fry, James, & Ferry, 2020; Müller, Vogel, Bai, & Vorholt, 2016; Zilber-Rosenberg & Rosenberg, 2008). These microbial communities contribute to the function of these organs: millions of years of intimate interactions between the host and its microbiome have forged pervasive interconnections between both parties. The microbiome plays a fundamental role in the development and function of the host immune system in both plants and animals. However, at the same time, the host immune system has been proposed to act as the resistant environment that imposes ecological filters on the microbial organisms, and thereby has the potential to shape host microbial communities (Hooper, Littman, & Macpherson, 2012; Lee & Mazmanian, 2010; Thaiss, Zmora, Levy, & Elinav, 2016). While the potential importance of the interactions between microbiome and immune system in determining the health of organisms has been studied in a few organisms, within-species diversity of microbiomes remains largely unstudied in non-human organisms (Bolnick et al., 2014; Garud & Pollard, 2020; Montero et al., 2021).

Major histocompatibility complex (MHC) genes encode proteins essential for the adaptive immune response of jawed vertebrates (Flajnik & Kasahara, 2001; Ohta et al., 2000). These molecules are essential for cell-mediated immunity and appeared early in the evolution of the adaptive immune system 500 million years ago (Flajnik & Kasahara, 2001; Rock, Reits, & Neefjes, 2016). The extensive population-level allelic diversity observed for these genes, alongside their central role in the vertebrate immune system, make them ideal candidates for studying how the host immune system affects its microbiota composition in wild populations. The study of this relationship in wild populations will also help understanding the reciprocal interplay between microbiota and the immune system shaping beneficial animal-microbial combinations, pathogen elimination, and disease resistance.

The influence of the MHC haplotype on the microbiome has been studied in all major vertebrate groups including fish (Bolnick et al., 2014), amphibians (Belasen et al., 2021; Hernández-Gómez, Briggler, & Williams, 2018), birds (Darolová, Poláček, Krištofík, Lukasch, & Hoi, 2021; Leclaire et al., 2019), and mammals (Khan et al., 2019; Kubinak et al., 2015; P. Lin et al., 2014). In humans, MHC (known as human leukocyte antigen, HLA) variants have been associated with the composition of the microbiome (Bolnick et al., 2014; Bonder et al., 2016). The results found in these studies offer three conflicting predictions on MHC-microbiota interactions. 1) A negative correlation between MHC diversity and heterozygosity, and microbiota diversity (Bolnick et al., 2014; Leclaire et al., 2019) where a higher diversity of MHC haplotypes leading to higher diversity of peptides presented to eliminate a higher number of microbe species. 2) A positive relationship where higher MHC diversity causes higher microbiome diversity (Hernández-Gómez et al., 2018; Khan et al., 2019), because not only do the immune system eliminates microbes, but also forms symbiotic bonds with commensals as higher diversity of MHC haplotypes initiating tolerance to a more diverse range of microbes. 3) MHC diversity is not related with microbiome diversity but its composition, where certain MHC haplotypes interact with specific groups of microbes, resulting in covariation between MHC genotypes and microbiome community structure (Bonder et al., 2016; Olivares et al., 2015). Note that statements 2 and 3 are not mutually exclusive.

Understanding the causal connections between the MHC and microbiota are especially relevant in groups where virulent wildlife diseases are contributing to population declines (Fisher et al., 2012). Among these diseases, chytridiomycosis stands out as an emerging disease caused by the chytrid fungi *Batrachochytrium dendrobatidis* (Bd) and *B. salamandrivorans* (Bsal) inflicting amphibian mass die-offs worldwide (Kilpatrick, Briggs, & Daszak, 2010; Martel et al., 2014; Scheele et al., 2019). Recent studies demonstrate the importance of the skin microbiota in amphibian innate immune defense against Bd (Bates et al., 2018; Rebollar et al., 2016; Torres-Sánchez & Longo, 2022). Hence, investigating how host MHC genetics, environment and evolutionary history determine the skin microbial diversity and composition of amphibian populations is a priority task in amphibian conservation (Jiménez & Sommer, 2017; Trevelline, Fontaine, Hartup, & Kohl, 2019).

Despite recent calls for an integration of microbiome research in evolutionary and conservation biology (Cullen et al., 2020; Henry, Bruijning, Forsberg, & Ayroles, 2021; West et al., 2019), little progress has been made on the fundamental association between host population history and genetic variation, and the diversity and composition of host microbiome in wild populations. Here, we studied the variation in MHC Class II in 12 moor frog *Rana arvalis* populations from Scandinavia originating from different environments and having different evolutionary histories. Previous studies have demonstrated that the postglacial colonization processes after the Last Glacial Maximum had a profound impact on the geographical distribution of *R. arvalis* and its genetic diversity (Cortázar-Chinarro et al., 2017; Knopp & Merilä, 2009; Rödin-Mörch et al., 2019). The results indicate that current patterns of MHC variation across Scandinavia reflect two different postglacial colonization routes and show signatures of past and ongoing differential selection patterns, drift, and historical demographic events, where southern populations have higher haplotype richness than the ones in the north (Cortázar-Chinarro et al., 2017; Cortazar-Chinarro, Meyer-Lucht, Laurila, & Höglund, 2018).

The inferred local adaptation in the moor frog is expected to not only be determined by the host genome, but also by the genetic repertoire of the microbiome, which, in turn, is affected by the major evolutionary forces of selection, drift, migration and mutation. Consequently, the host can be expected to be under strong selection to shape the microbiota and foster a beneficial microbial community (Foster, Schluter, Coyte, & Rakoff-Nahoum, 2017). Investigating this relationship in an evolutionary context is imperative in order to understand the distribution of host-microbiome biodiversity, its evolutionary association history and the forces that have generated it (Groussin, Mazel, & Alm, 2020).

Here, we ask the following questions: (i) Does geography and/or host evolutionary history affect the diversity and structure of the skin microbiota? (ii) Is MHC heterozygosity correlated with microbial diversity? (iii) How does MHC diversity affect microbiome? and (iv) Does MHC haplotype similarity correlate with microbial diversity and/or skin microbiota composition?

## Methods

### Study sites and sampling

*Rana arvalis* has a broad longitudinal and latitudinal distribution in Eurasia and is relatively common in most of Fennoscandia (Wielstra, Sillero, Vörös, & Arntzen, 2014). Previous studies showed a bidirectional postglacial colonization route of the species to Scandinavia: a western lineage coming from the south via Denmark to Southern Sweden and another lineage arriving from the east via Finland to northern Sweden (Cortázar-Chinarro et al., 2017; Knopp & Merilä, 2009; Rödin-Mörch et al., 2019). Eight sites close to Uppsala (Uppland region, henceforward termed ‘South’) corresponding to the western lineage and four sites in Luleå (Norrbotten region, henceforth called ‘North’) corresponding to the eastern lineage were selected as sampling locations in this study. Study sites within each region were at least 8 km apart and differing in habitat (i.e., from open farm ponds to forest ponds). Sampling was conducted during the breeding season in March – April (South) and May (North) 2016 (Table S1).

A total of 207 adult frogs were captured using hand nets. Each individual was handled with a new pair of sterile nitrile gloves to avoid cross contamination. All individuals were sexed and weighed prior to sample collection. Sample collection included removal of a piece of tissue from the toe webbing and storing it in 90% alcohol for DNA extraction. To sample the skin microbiome, each frog was transferred to an individual 250 mL container containing sterile distilled water (the Millipore Milli-Q™; Fisher Scientific) to remove transient microbes from the environment. After five minutes, each animal was moved to a new container with sterile distilled water and kept there for another two minutes. Finally, each frog was manually cleansed (again with sterile distilled water; ddH_2_O). The frogs were then carefully swabbed with a sterile rayon tipped MW100 (mwe; medical wire & Equipment Co) six times on both dorsal and ventral surfaces, covering as much skin as possible. Swabs were transported on ice in cooler boxes prior to storage at −80 °C in the laboratory.

To control for environmental microbes that might be found on the frogs’ skin, a 2L-water sample was taken from every study site. Water samples were taken within close proximity to where the frogs were captured from using a sterilized Durham glass bottle. Samples were kept cold and dark until processed in the laboratory. Water samples were filtered under a sterilized hood in the laboratory in the night of collection. As a pre-filtration step, two blank filtered samples (FNC1 and FNC2) were obtained from every water sample after filtering 200 mL DNA/RNA-free Milli-Q water. Bacterioplankton cells were collected onto 0.2 µm membrane filters (Super-200 Membrane Disc Filters; Pall Corporation, East Hills, USA), filtering 0.2 L of pre-filtered (0.7 µm; membrane filter) water. Pre-filtration was carried out to avoid capturing larger particles. Four water samples were taken at each site. Filters were kept at −80 °C until DNA extraction.

The temperature of every pond was recorded at the day of sampling using a portable multiparameter meter. Monthly temperature and precipitation (Worldclim data base: http://www.worldclim.org average of 30 years) at each sampling location were extracted to estimate the average values of these bioclimatic variables from the beginning of the breeding season in March to the end of the growing season in October.

### DNA extraction and Illumina MiSeq library preparation and sequencing

#### MHC Class II exon 2

The DNA from the tissue was extracted by using the DNeasy Blood and Tissue kit (Qiagen, Sollentuna, Sweden) following the manufacturer’s instructions. The complete second exon (272 bp) of the single MHC II gene (corresponding to the β −2 domain) in *R. arvalis* was amplified using the primers ELF_1 (3′-GAGGTGATCCCTCCAGTCAGT-5′) and ELR_2 (3′-GCATAGCAGACGGAGGAGTC-5) (Cortázar-Chinarro et al., 2017). Both forward and reverse primers were modified for Illumina MiSeq sequencing with an individual 8 bp barcode and a “NNN” sequence (to facilitate cluster identification). PCR reactions and library preparation are described in detail in Cortazar-Chinarro et al. (2017). A total of six libraries were generated using the ThruPLEX DNA-seq 6S (12) kit (Takara Bio Europe, TOWN, France). The concentration of each sample pool was measured with Quant-iT PicoGreen dsDNA assay kit (Invitrogen Life Technologies, Stockholm, Sweden) on a fluorescence microplate reader (Ultra 384; Tecan Group Ltd., Männedorf, Switzerland). The six libraries were combined in equimolecular amount of each sample into a MiSeq run, prior to sequencing. Sequencing of two MiSeq 250 (rxn) runs were carried out at the NGI/SciLifeLab Uppsala (Sweden).

#### Bacterial DNA extraction and library construction

The whole community DNA was extracted from both the swabs and filters using the DNeasy PowerSoil kit (Qiagen) following the manufacturer’s recommendations. Extracted DNA was sized and quantified by means of agarose (1.5 %) gel electrophoresis, GreenGel staining (Biotium Inc., Hayward, USA), and safe blue light transillumination prior to PCR amplification.

The bacterial swab and lake samples were subjected to 16S rRNA gene amplicon sequencing on an Illumina MiSeq platform (Illumina Inc., San Diego, USA). The sequencing library was prepared according to a two-step PCR. The first PCR step (30 cycles) amplified the bacterial hypervariable region V4 of the16S rDNA gene, using bacterial forward primer 515F (5′-GTGCCAGCMGCCGCGGTAA -3′) and reverse primer 806R (5′-GGACTACHVGGGTWTCTAAT -3′) (Varg et al., 2022). The second PCR step (20 cycles) attached indices to both ends of the 16S amplicons in order to create a unique dual barcode for each individual sample (See Table S2 and additional information A1). The 16S primers used in the first PCR step were, thus, modified by means of extending their 5’-ends with Illumina adapter sequences, These barcoding primers also comprised the Illumina sequencing handle sequence, which attaches the amplicons onto the Illumina flow cell to initiate sequencing. In both PCR steps, the Phusion high-fidelity DNA polymerase (ThermoFisher Scientific, Stockholm, Sweden) was used, and PCR mixtures were each time prepared according to the manufacturer’s instructions with the addition of 20mg/mL of BSA (Bovine Serum Albumina, Thermo fisher, Stockholm, Sweden). Also, amplicons were purified after each PCR step using the Agencourt AMPure XP purification kit (Beckman Coulter Inc., Brea, USA), and amplicon fragment size and quantification were checked using a Bioanalyzer (Agilent) and a fluorescence microplate reader (Ultra 384; Tecan Group Ltd., Männedorf, Switzerland), employing the Quant-iT PicoGreen dsDNA quantification kit (Invitrogen). Finally, equimolar amounts of samples were mixed, and the final amplicon sequenced using the Illumina MiSeq platform (Illumina Inc.) at NGI/SciLifeLab Uppsala (Sweden).

#### MHC Class II exon 2 sequence processing

Sequence processing was performed following (Cortázar-Chinarro et al., 2017). The raw amplicon sequencing was combined into single forward reads using FLASH (Magoč & Salzberg, 2011). Each of the six amplicon pools was analyzed independently. A total of 6 fastq files were generated and transformed to fasta by using Avalanche NextGen package (DNA Baser Sequence Assembler v4 (2013), Heracle BioSoft, www.DnaBaser.com). AmpliCHECK (Sebastian, Herdegen, Migalska, & Radwan, 2016) was used for removal of primer sequences, de-multiplexing, chimera removal and counting variants for each amplicon, while AmpliSAS (Sebastian et al., 2016) was used for final allele verification. The DOC method (Lighten, Van Oosterhout, Paterson, McMullan, & Bentzen, 2014) was used, where variants are sorted top-down by coverage, followed by the calculation of the coverage break point (DOC statistic) around each variant. Individuals in which replicate samples due to PCR artefacts that did not reveal identical genotypes were removed from the consecutive analyses. We retained samples that were reveling an identical genotype for at least two out of three replicates. All valid allele sequences from 179 retained individuals were imported and aligned in MEGA X (Kumar, Stecher, Li, Knyaz, & Tamura, 2018). All sequences were extensively compared to other sequences from the same locus (*R. arvalis*: GenBank: isolates from h1 to h57 -MT002608.1-MT002664.1]). We used the MHC nomenclature by (Klein 1975) for the valid retained alleles. This nomenclature consists of a four-digit abbreviation of the species name followed by gene*numeration, e.g. Raar_DAB*01.

#### Bioinformatic processing of bacterial data

Raw sequences were processed using DADA2 (Callahan et al., 2016). Forward and reverse reads were trimmed to 240 and 200 bp, respectively, using default parameters. Default parameters were, moreover, employed to correct for amplicon errors, identify chimeras, and merge-end reads. Taxonomic assignment following amplicon sequence variant approach (ASV) was performed with the help of the bacterial 16S rRNA SILVA reference data base (version v132) training set (Yilmaz et al., 2014). All unassigned ASVs were removed from the samples (Costa, Tavares, Baptista, & Lino-Neto, 2022; Couch et al., 2021). Furthermore, to minimize erroneous ASVs, all singletons were removed according to the default settings of DADA2. The data was filtered by sample or taxa, using the functions *subet_sample*, *prune_taxa* () implemented in the Phyloseq R package. For statistical analyses, data was transformed into proportions (compositional data) in order to minimize erroneous ASVs and for direct count comparisons which is a modern data that do not need rarefraction to produce correct results (Cameron, Schmidt, Tremblay, Emelko, & Müller, 2020; McMurdie & Holmes, 2014; Willis, 2019). We used used the function *transform_sample_counts (ps, function (out) out/sum(out))* implemented in the Phyloseq package (McMurdie & Holmes, 2013) or with package Microbiome (Shetty & Lahti, 2019) in R. We used the “*compositional*” method and “*clr*” (i.e. relative abundance methods). The analyses were verified with all the methods.

#### MHC Class II exon 2 data analyses

We assessed genetic diversity in MHC Class II exon 2 using standard diversity indices (HE, HO, allelic frequencies, nucleotide diversity). These were calculated for each locality in Arlequin v 3.5 (Excoffier & Lischer, 2010). Allelic richness was calculated in FSTAT 2.9.3.2 (Goudet, 1995); See Table S3). Allele frequency plots were created in R using the “ggplot2” package (Wickham, 2016).

To collapse MHC alleles into functional supertypes, we extracted the 12 codon positions for the Peptide Binding Region (PBR) according to (Cortazar-Chinarro et al., 2018). We then characterized each codon based on five physiochemical descriptor variables: z1 (hydrophobicity), z2 (steric bulk), z3 (polarity), z4 and z5 (electronic effects) (Jombart, Devillard, & Balloux, 2010). A hierarchical clustering tree for the MHC class II exon 2 in *R. arvalis* was constructed with the z-descriptors in R (version 4.0.5). The optimal number of clusters was decided based on divergence between the branches in the phylogenetic tree. Alleles within each cluster was collapsed into a single Supertype (Figure 1). Supertype allele frequency plots were created in Excel -See Figure 1]. Note that even if we consider supertyping as useful method for investigating broad associations between MHC, microbiome and infection in field studies, we recognize the limitations of using supertypes in our study, as recent studies have demonstrated that the size of peptide repertoires is not correlated with peptide motifs of many MHC class I molecules supertypes. These results call the immunological relevance of supertypes into question (Kaufman, 2020; Tregaskes & Kaufman, 2021).

**Figure 1.**
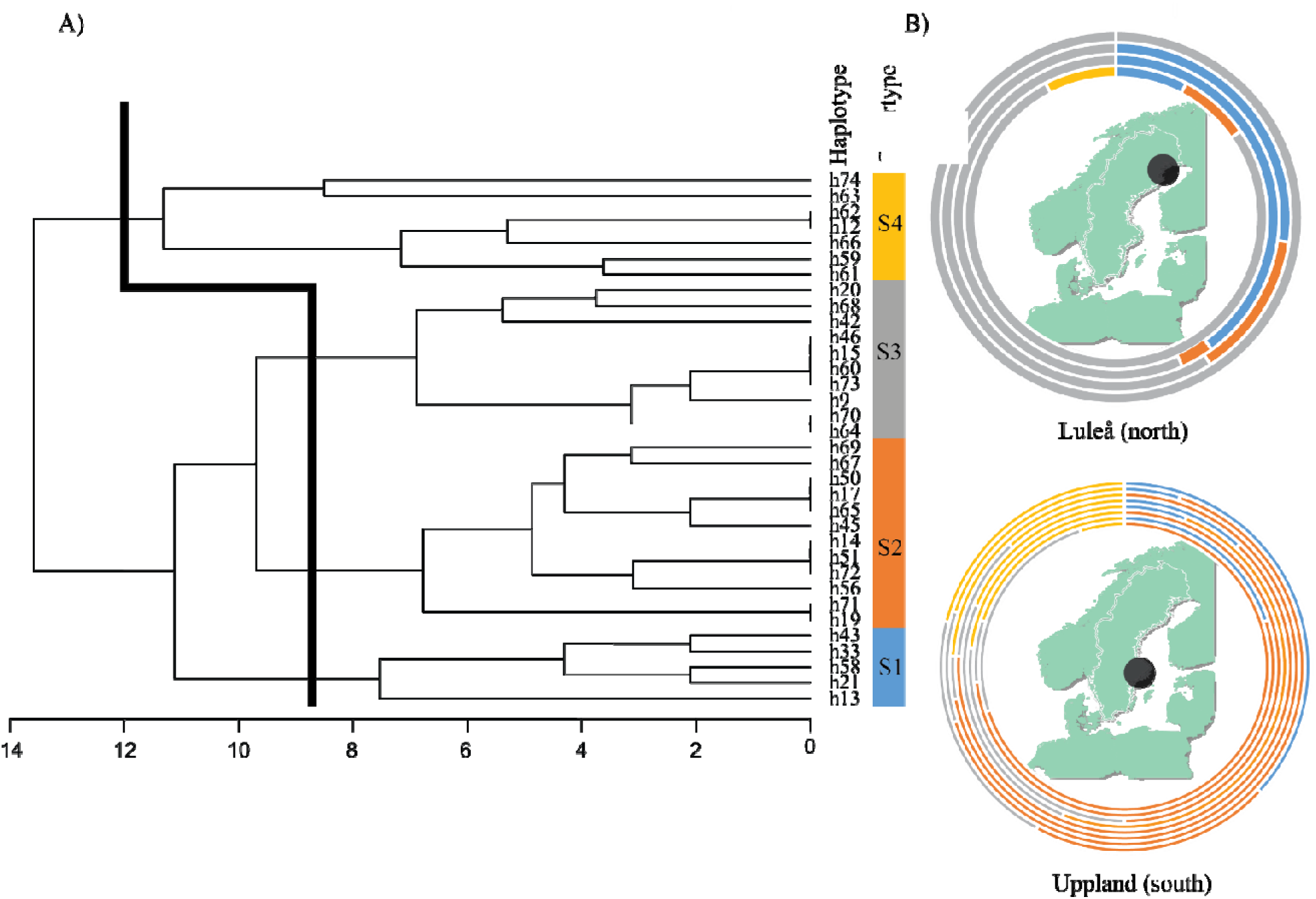
A) A hierarchical clustering tree based on the Peptide binding regions (PBR) of MHC class II exon 2 in *R. arvalis and* the z-descriptors recovering a total of 34 haplotypes. The black line indicates the optimal number of clusters grouped as supertypes (S1, S2, S3 and S4). The number of clusters (supertypes) was based on the divergence between the branches in the phylogenetic tree. Alleles within clusters were collapsed into a single supertype. B) Pie chart representing supertype frequencies in the two regions (Luleå (north) and Uppland (south) marked with a black dot).

#### Bacterial diversity data analyses

Differences in composition between the environmental pond and amphibian skin communities were examined by means of permutational multivariate analysis of variance (PERMANOVA, analysis of differences in group means based on distances, 999 mutations) and permutational multivariate analysis of dispersion (PERMDISP), analysis of differences in group homogeneities based on distances.

Bacterial alpha-diversity was estimated using Observed richness, Shannon diversity, and phylogenetic diversity indices. Comparisons between regions and sexes were carried out using Wilcoxon and Kruskal-Wallis tests implemented in Phyloseq (McMurdie & Holmes, 2013) and Picante (Kembel & Kembel, 2014) R packages. Correlation coefficients between ASVs Observed richness and total number of reads were also assessed to demonstrate that the asymptote of the samples has been reached and thus no misinterpretation of the diversity is occurring (Figure S1). GLM and GLMMs with Gaussian error structure were used to assess whether alpha-diversity (Shannon) could be explained by the environmental factor: 1) Temperature at time of sample collection “TemCollection”, 2) Average temperature “TemMean”, and 3) Average precipitation “PreMean” on ASVs bacterial diversity. Population was included as a random factor.GLMMs models were run in R (Team, 2013).

Differences in bacterial composition among communities between the two regions (South and North) and sexes were analyzed using PERMANOVA by using *Adonis2()* function in Vegan package with 999 permutations and test of homogeneity of group dispersion (PERMDISP) on weighted and unweighted UniFrac distances. Relationships between the bacterial assemblages from the South and North were explored employing hierarchal cluster analyses (Bray-Curtis distance and UniFrac distances by using Vegan (Oksanen et al., 2019) and circlize (Gu, Gu, Eils, Schlesner, & Brors, 2014) packages implemented in R (Team, 2013).

#### Associations between bacterial diversity and host MHC class II exon 2

Relationships between MHC genetic and bacterial diversity (Shannon, Simpson and Chaos1) were analyzed by several tests. Shannon diversity index was the only diversity index that was perfectly adjusted with the structure of our data. Therefore, the analyses in the following only consider the Shannon diversity index. First, we assessed the effect of MHC heterozygosity and bacterial Shannon diversity at the population level. Second, we investigated the effect of MHC nucleotide diversity and bacterial Shannon diversity at the individual level. For both, multiple regression on distance matrices (MRM) and (lm) models were used with the package Ecodist (Goslee & Urban, 2007) and nlme in R (Team, 2013).

Relationships between heterozygosity at the supertype level and Shannon bacterial diversity index between regions were explored by performing a GLMM model in R. Individuals were, for this purpose, grouped in two categories: “*sameS*” and “*DistinctS*”. *SameS* individuals were defined as individuals in which the two alleles belonged to the same supertype group (e.g., Supertype2_2), while *DistinctS* individuals carried two alleles that belong to different supertype groups (e.g., Supertype1_3). We assume that heterozygosity is lower within the “*SameS”* group compared to the “*DistinctS”* group, as the probability of having the same two alleles is higher within “*SameS”* group. Individual Shannon diversity index was used as the response variable and region as a fixed factor with population as a random effect. Additional GLMM analyses were carried out to test for differences between bacterial diversity and specific supertype-haplotype groups within the regions. Moreover, redundancy analyses (RDA) in the Vegan package (Oksanen et al., 2019) were run to find potential indications of a relationship between bacterial community composition and supertype haplotype structure. Likewise, we used RDA to summarize linear relationship between the bacterial community composition and MHC specific supertypes.

DESeq2 and ANCOMBC2 was performed to explore whether specific bacterial taxa differed in abundance between “*SameS*” and “*DistinctS*” groups of individuals as well as between specific supertypes (H. Lin, Peddada, & Lin, 2021; Love, Anders, & Huber, 2014). In addition, ASV abundance and supertype data were cross-correlated to find specific taxa correlated with specific supertypes (Spearman’s rank correlations). First, we transformed abundance data into compositional data by using Microbiome package (Shetty & Lahti, 2019) in R software. The neighbor-joining tree, showing the phylogenetic relationships among ASVs negatively and positively correlated to MHC supertypes was constructed with MEGA X (Kumar et al., 2018).

## Results

### MHC II exon 2 and Skin Micriobiome characterization

We obtained a total of 4.2 million reads with intact primers and attached barcode information that could be assigned to 207 individuals. We amplified and sequenced in duplicates or triplicates, which corresponds to 81.8% of the total number of samples. One sample out of 321 failed due to PCR amplification problems. The average number of reads per amplicon was 13085.37 ranging from 420 to 106172 reads, 2.7 million reads remained after filtering and QC analyes. We assigned 34 valid MHC class II exon 2 alleles with a length of 272 bp and 27 polymorphic nucleotide positions among the 179 remaining individuals. All the 34 valid MHC II exon 2 allele sequences were translated into unique amino acid alleles. 17 out of the 34 alleles were found in a previous study (Cortázar-Chinarro et al., 2017) and another 17 were new alleles discovered in the present study (Raar_58 to Raar_74). By following the DOC method (Lighten et al., 2014), we detected a single locus in 193 individuals. Three individuals showed evidence of a second MHC class II locus with apparently lower number or reads in two of the three replicates, pointing to the possible existence of a very rare MHC class II duplication. We conclude that we are mostly working with a single MHC class II locus in our data set. However, we cannot rule out the possibility our primers amplify an MHC class II locus in a few cases (two individuals).

A total of 37148 reads were obtained from both amphibian (n=179) and water samples (n=12), with amphibian swabs contributing 84.84% to the total number of reads. The most abundant phyla were Proteobacteria (45% of the total number of sequences), Bacteroidetes (16%), Actinobacteria (9.9%), Acidobacteria (6.77%), Verrucomicrobia (5.39%), and Firmicutes (3.71%). The rest of the phyla represents less than 2.5% of the total number of reads: Planctomycetes (2.1%), Chloroflexi (1.77%), Armatimonadetes (1.4%), Candidatus_ Saccharibacteria (0.81), and Gemmatimonadetes (0.75%) (Figure S2). After removal of uncharacterized taxa (n=1280 ASVs; 7.8% of the total abundance), 15017 taxa remained.

### Genetic diversity

#### MHC class II exon 2

The number of alleles per population varied substantially between the northern and southern region (Figures 2, S3 and Table S3). Levels of expected heterozygosity for the MHC locus between populations ranged from 0.23 to 0.84 (Overall H_E_=0.79, Table S3) and allelic richness ranged from three to 11 (overall AR=7.83, Table S3). The northern region showed lower diversity than the southern region in terms of H_E_ and AR. Two alleles were private to a single population in the southern region (Raar_69 and Raar_65; Figure 2, S2). Three alleles were only present in the northern region at low frequencies and private to a single population (Raar_42, Raar_43, Raar _68). However, Raar_42 and Raar_43 were found in the southern region in a previous study -33] (Figure 2, S3).

**Figure 2.**
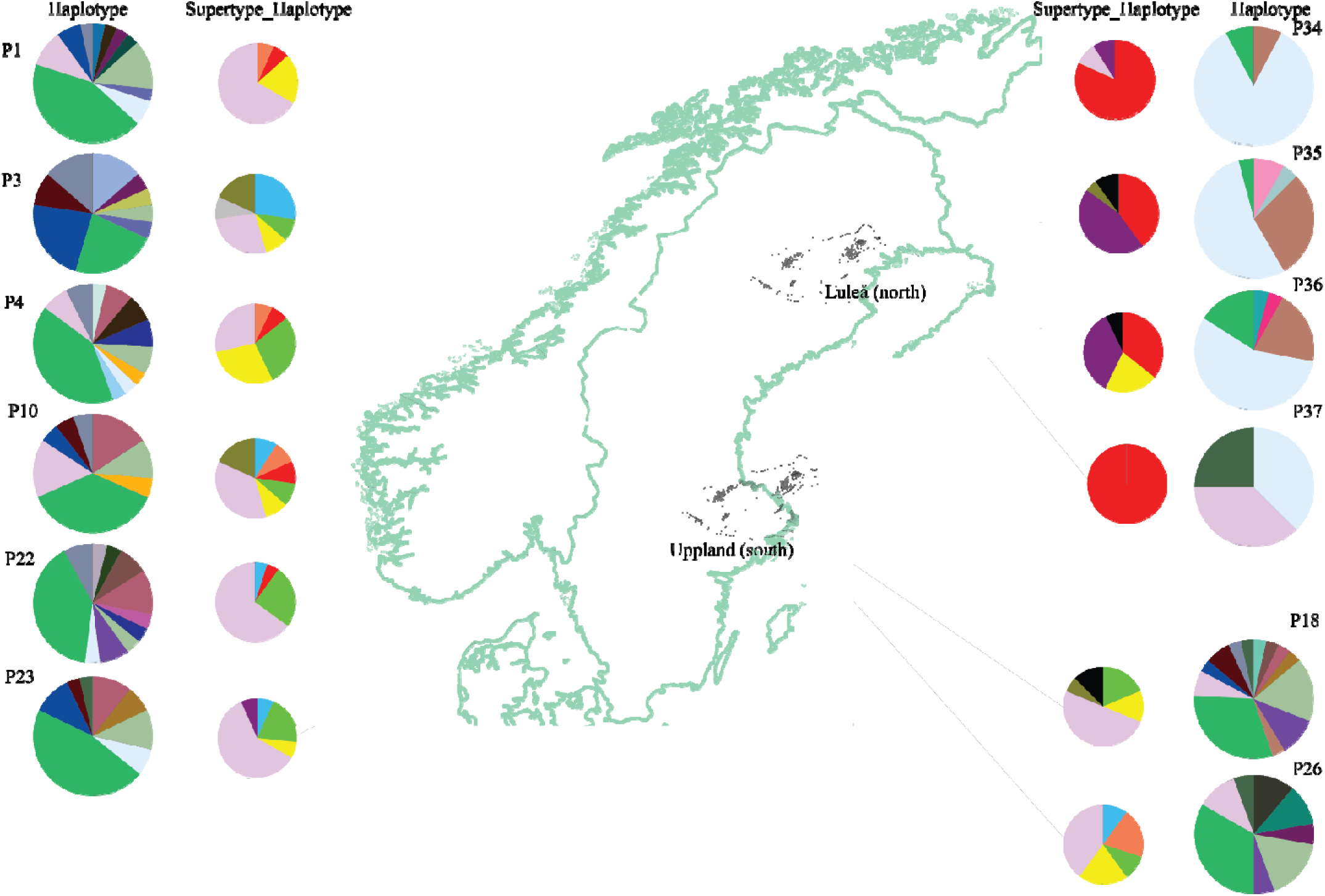
Allelic frequency distribution of MHC Class II haplotypes and supertype haplotypes in 12 *R.arvalis* populations (P1: Ekeborg, P3: Eneby, P4: Valsbrunna, P10: Kroklösa, P22: Högbyhatt, P23: Dalkarlskärret, P18: Mosta, P26: Ströbykärret (Uppland), P34: Lillträsket, P35: Vittjärnen, P36: Djurhustjärnen, P37: Dalbacka (Luleå). Colour coding scheme for MHC alleles is given in Fig. S2.

The 34 alleles were converted into four different MHC class II exon 2 supertypes based on physiochemical binding properties (Figure 1). Supertype_2 was the most common supertype in the southern region while Supertype_3 was the most abundant in the northern region (Figure 1). Supertypes were also grouped by genotypes. Supertype_Haplotpe_ diversity was defined as the diversity within each genotype. Supertype_Haplotype_ was higher in the southern region (Supertype_Haplotype_south_=9; Supertypes_Haplotype_north_=6). 49% of the southern individuals carried the Supertype_Haplotype_2_2 while only 1.18% carried the Supertype_Haplotype_2_2 in the north. By contrast, Supertype_Haplotype_3_3 and Supertype_Haplotype_1_3 occurred at higher frequency in the north than in the south (3_3: 58.1% and 27.2% and 1_3: 3.53% and 0.88%, respectively) (Figure 2; S3).

#### Skin bacterial diversity patterns in relation to environmental variables, regions, and sex

We found significant differences between bacterial community composition in the water filters and on the skin of the amphibians (PERMANOVA; p<0.05, PERMDISP); p<0.05; See Figure S4, Table S4). Alpha-diversity was similar for both sexes (Wilcoxon Observed; W = 3486, p = 0.33; Wilcoxon Shannon; W = 3453, p = 0.3947; Wilcoxon PD; W = 3469, p = 0.36) and regions (Wilcoxon Observed; W = 2901.5, p = 0.8895; Wilcoxon Shannon; W = 2954, p = 0.9663; Wilcoxon PD; W = 2959, p = 0.95; See Figure S5). Average temperature (PreMean) and Temperature at data collection (TemCollection) and for “Region” were positively related to alpha-diversity (See Table S5). When we controlled by population, the average precipitation (PreMean) was significantly different depending on the region, indicating high heterogeneity in alpha diversity with precipitation and differed by origin. However, we did not observe significant effects of the average temperature or the temperature at collection time (TemCollection) on the skin microbiota diversity on GLMMs models (Table S5). Additionally, we found support for a regional effect in Beta-diversity, showing significant differences in bacterial community composition between the two regions (PERMANOVA; p<0.05 weighted and unweighted UniFrac distances, Figure 3; S6 and Table S6) but not differences in group dispersions (PERMDISP; p>0.05, see Figure S7, and Table S7).

**Figure 3.**
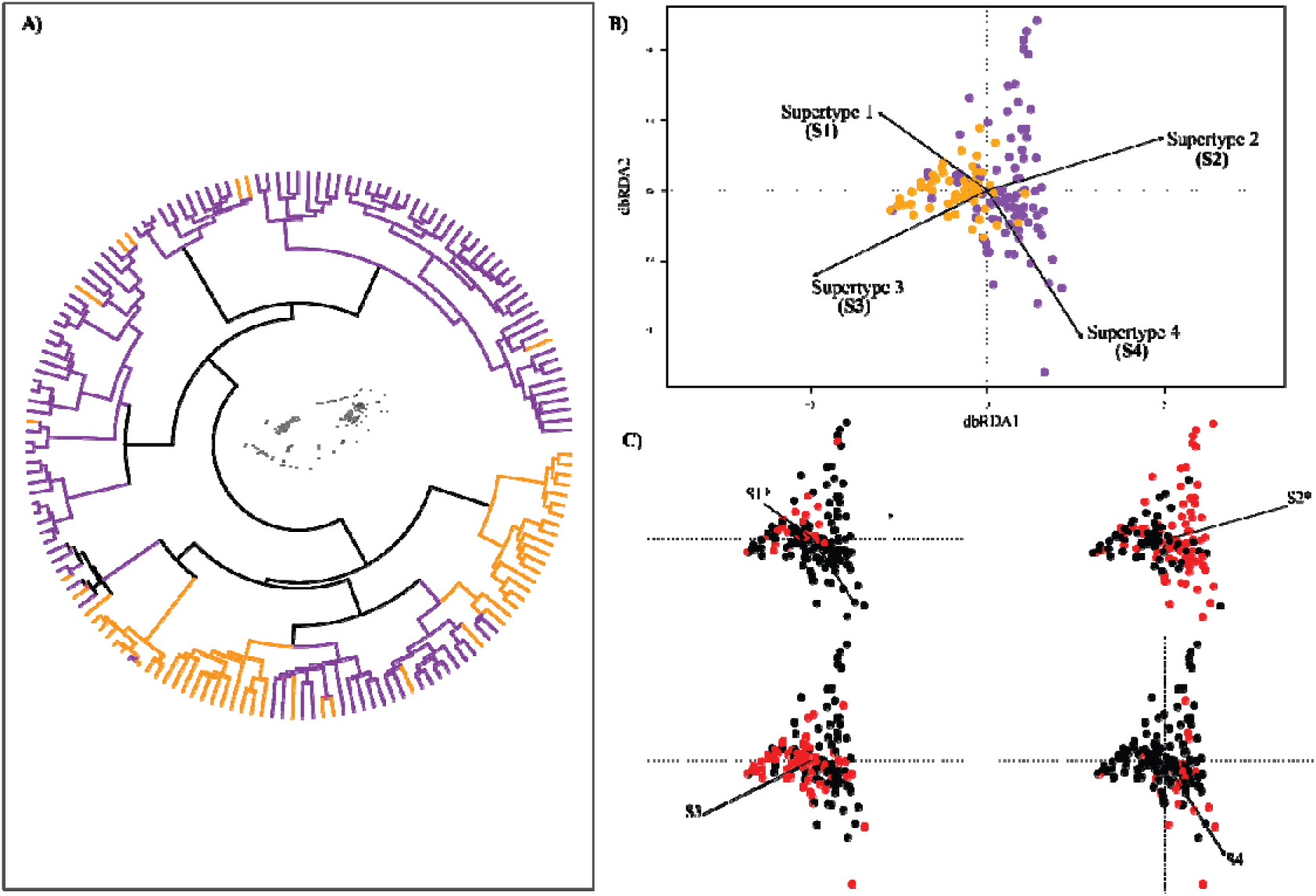
**A)** Differences in bacterial community composition of 16S DNA skin microbiota between regions represented by hierarchal clustering of samples (Ward′s clustering; Bray-Curtis distance). Clusters representing the 16S DNA skin microbiota composition from Uppland (South) are colored in purple and from Luleå (North) in orange, respectively. **B)** RDA performed with the bacteria identified in skin microbiome clustered in two main groups according to the amphibian origin. Each point represents the skin microbial community of an individual *R. arvalis*. 16S skin samples of an individuals from Uppland are represented in purple and samples of individuals from Luleå are represented in orange. The supertypes (S1, S2, S3 and S4) are settled as variables represented with arrows. **C)** The RDA plots show the separation pattern for every single supertype (S1, S2, S3 and S4). The supertypes that are represented in plot B and C are highlighted red and labelled in bold and with an increase in size. Significant supertypes are marked with (*) with a p<0.05 according to PERMANOVA, Adonis test.

### Effects of MHC class II heterozygosity and Skin microbiota composition

At the population level, populations with higher MHC heterozygosity exhibited more diverse microbiota according to the multiple regression analyses (See Table S8 and Figure S8). However, individuals with more divergent MHC sequences based on the nucleotide diversity had less diverse microbiota (Figure S8, Table S8). Given these results, we infer that protein structure dissimilarity among MHC sequences within a host reduces the diversity of skin microbial communities based on the nucleotide diversity values. Nucleotide diversity is directly correlated to the *Theta k* value, which is a proxy of MHC sequence dissimilarity. Table S3 show that *theta k* (k) values clearly differ between populations and regions, being lower at northern region where MHC sequences are more similar one to each other (See Figure S9, Table S8).

We next quantified the relationship between individuals that carried MHC alleles grouped within the same supertype cluster “homozygous” and individuals that carried MHC class II alleles. The alleles were grouped in different supertype clusters “*heterozygous*” individuals in respect to the overall diversity. We grouped all the “*homozygous*” and “*heterozygous*” individuals according to their supertype clustering information. Using GLM models, we found that “*DistinctS*” individuals from the north showed a significantly higher microbial diversity in comparison to southern *“DistinctS”* individuals (Figure S10 and Table S9). However, we did not observe such effect on the *“SameS”* individuals, suggesting a potential bacterial diversity compensation for the deficiency of MHC diversity in the northern region. We did not find significant differences in bacterial diversity between specific MHC genotypes present in both regions (Supertype 2_3 and Supertype 3_3). Furthermore, RDA and PERMANOVA analyses did not show beta bacterial diversity patterns between “*DistinctS*” individual group (MHC class II alleles grouped in different Supertypes) and “*SameS*” individuals group (MHC class II alleles grouped in the same Supertype). However, our data show significant differences in community structure among specific supertype-haplotypes (See Table S10). Additionally, we found that individuals carrying supertypes 1 or 2 had a specific bacterial composition (See Figure 3B and 3C; Figure S10 and Table S11). Individuals carrying Supertype haplotype 1_3, 2_2, 2_4 or 3_3 present a specific bacterial composition (See Table S12). Both these results might be explained as a strong direct regional effect between north and south as well as an effect of a specific combination of MHC class II exon 2 on the microbial structure (Figure S11).

### Associations between MHC supertypes and microbial taxa

We did not find bacterial ASVs that were significantly different in abundance between “*SameS*” and “*DistincS*” individuals according to the MHC supertype clustering. On the contrary, we found bacterial ASVs that were significant in abundance per supertype (See Table S13 S14; Figure S12). *Oxalobacteraceae* family was the most common taxa found by using both DESEq2 and ANCOMBC2. Likewise, the heatmap (Figure 4B) illustrated positive and negative correlations (spearman rank correlation p<0.05) between supertypes and specific ASVs. Families exhibiting significant MHC effects (*Comamonadaceae, Oxalobacteraceae and Pseudomonadaceae*) are taxonomically clustered (Figure 4A). Supertype 4 affects the abundance of one single family of *Bacteroidetes*, whereas supertypes 1, 2 and 3 affect the abundance of at least two *Proteobacteria* families. Surprisingly, Supertypes 2 and 3 which are the most dominant supertypes in the southern and northern regions, respectively, show an antagonistic association in specific bacterial taxa, especially *Oxalobacteraceae*.

**Figure 4.**
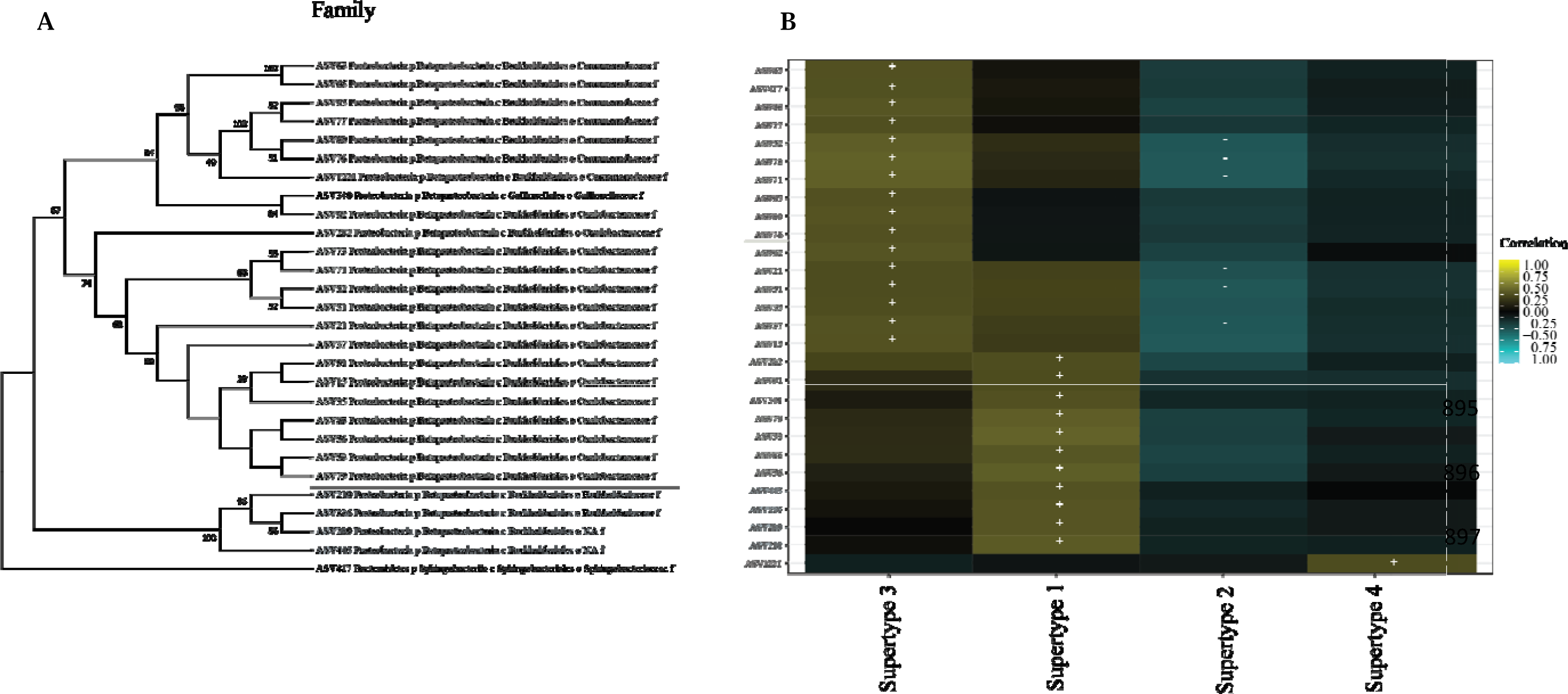
**A)** The neighbor-joining tree shows the phylogenetic relationships among ASVs correlated to MHC supertypes at family level. *Commamonadaceaeae* family is colored in dark pink, *Oxalobacteraceae* is colored in light green and bacteria belonging to the *Burkholderiaceae* family are colored in light blue. **B)** Heatmap showing the correlations between supertypes and specific ASVs. Significant spearman cross-correlations (p <0.05) are labeled with (+) or (-). Positive correlations are shown in dark yellow (+) and negative correlations (-) in dark

## Discussion

We characterized the skin microbiota composition and MHC Class II exon 2 diversity in 12 *R. arvalis* populations from two separate geographical regions representing different evolutionary histories due to different post-glacial colonization histories. We also assessed the relationships between MHC genotype and microbial community diversity to investigate potential associations between the host MHC genes and skin microbiome and to elucidate differences between regions and evolutionary histories. Our results indicate that the skin microbial community of frog populations varies substantially among populations and regions. The climatic and pond environmental factors appeared to influence the diversity and structure of microbial communities, but most of the differences identified could not be explained by environmental factors in our study. Therefore, influence of the microbial community structure could be related to the host genetic variation, although our data cannot prove such relationship. Further investigation should be carried out in this regards. Four main results can be derived from our analyses. First, alpha diversity was similar between regions while beta diversity, which is related to microbial composition, was significantly different between the regions. Second, within populations, MHC heterozygosity was positively correlated with microbial alpha diversity. Third, heterozygous individuals from the north showed higher alpha diversity compared to the heterozygous individuals from the southern region, where MHC class II diversity was higher. Fourth, there were indications of antagonistic associations between MHC class II alleles and specific bacterial taxa at the regional level. We will discuss each of these results in detail below.

### Microbiome variation between regions and populations

Previous studies have shown that genetic distribution of MHC class II alleles was strongly influenced by evolutionary processes such as migration, drift, selection, and demography in amphibians (Cortázar-Chinarro et al., 2017; Luquet et al., 2019). However, very little is known about how evolutionary processes influence skin microbiota diversity in amphibians (Belasen et al., 2021; Torres-Sánchez & Longo, 2022). Despite that our results suggested a similar pattern of alpha diversity between regions and populations, the relative abundance of shared ASVs, beta diversity and, thereby, the bacterial community structure composition varied between regions and populations. Regions and populations had distinct skin microbial communities, likely reflecting differential environmental and host-specific filtering, where historical genetic background of different colonizing lineages as well selective pressures may have an important role of host-microbiome biodiversity distribution. The effect of the genetic background of the host has been proposed as a stronger predictor of skin microbiome structure in other systems (Amato et al., 2016; Dimitriu et al., 2019; Muletz Wolz, Yarwood, Campbell Grant, Fleischer, & Lips, 2018; Weinstein et al., 2021). Earlier, it was found that co-occurring Panamanian frog species host unique skin bacterial communities (Belden et al., 2015). Despite this, it is unknown if host-associated traits, such as the immune genes, select for specific host bacterial communities in amphibians, as it does in other organismal groups such as humans (Shafquat, Joice, Simmons, & Huttenhower, 2014; Wein & Sorek, 2022). However, we cannot rule out the possibility that such host-pathogen associations might also be driven by linkage with other genes.

### Deterministic factors contributing to microbiome variation

Adaptive immune genes such as MHC have extensively been linked to the susceptibility to infections in vertebrates (Savage, Muletz-Wolz, Campbell Grant, Fleischer, & Mulder, 2019; Savage & Zamudio, 2011). Parasite-specific immune responses driven by MHC polymorphism have been studied to a great extent (Eizaguirre & Lenz, 2010; Elbers & Taylor, 2016; Minias, Whittingham, & Dunn, 2017; Peng, Ballare, Hollis Woodard, den Haan, & Bolnick, 2021). However, how the complex relationship between MHC and a multitude of host-associated microbes influence the host immune response is still poorly understood. While mammalian studies have highlighted that host genetic background can influence microbial communities via the immune system (Blekhman et al., 2015; Tabrett & Horton, 2020; Woodhams et al., 2020), less is known for other taxa. However, recent investigations have shed light on important associations between host immunity and microbiomes (Bolnick et al., 2014; Fleischer, Risely, Hoeck, Keller, & Sommer, 2020; Hernández-Gómez et al., 2018), not only for MHC genes themselves, but also for other immune and cell signaling genes linked to MHC Class I and II (Flajnik, 2018; Grogan et al., 2018; Richmond, Savage, Zamudio, & Rosenblum, 2009). In sticklebacks, high MHC variation has been associated with a diverse microbiota (Bolnick et al., 2014), while in amphibians high MHC variability may influence host health indirectly by shaping bacterial communities (Belasen et al., 2021). In concordance to previous findings in *R. arvalis* (Cortázar-Chinarro et al., 2017), we found lower MHC class II diversity at northern latitudes, conferring a possible increase in susceptibility to infection. However, we did not find regional differences at bacterial alpha diversity, but at the microbial community composition.

We found a positive link between expected MHC heterozygosity and bacterial alpha diversity. These results are in line with the heterozygote advantage (overdominance) theory where heterozygous individuals might successfully carry a highly diverse bacterial community on the skin and consequently heighten resistance to infection (Khan et al., 2019). Additionally, we found that more divergent MHC alleles are negatively associated to alpha diversity and heterozygous individuals from the northern populations carry a more diverse bacterial community as compared to individuals from the southern populations. This result suggests that lower genetic variation commonly observed at northern latitudes could be compensated by higher bacterial richness at northern populations, showing support to the idea that more diverse bacterial communities will compensate for the lower individual MHC diversity at northern latitudes.

Studies on chimpanzees have shown a straightforward relationship between a healthy and diverse immune system and the gut microbiome composition and the direct role on its body internal regulation (Barbian et al., 2018; Björk, Dasari, Grieneisen, & Archie, 2019). Consequently, individuals suffering immunodeficiency due to a pathogenic infection experience substantial alterations of their gut microbiota communities (Dillon et al., 2014; Moeller et al., 2015), confirming that microbes shape immune responses (Salas & Chang, 2014). In humans, patients with a poor immune system show higher gut bacterial diversity and higher frequency of low genetic diversity genes than patients with a regular immune system in terms of diversity (Bosák et al., 2021). Most of the studies have focused on gut microbiome and its direct association with the mammalian immune system and very little has been done on other bacterial communities (incl. skin microbiota) and other host vertebrate groups. Therefore, experimental studies investigating the role of skin microbiome diversity in shaping immune response are urgently needed in order to gain a better understanding of the factors causing the bacterial community compensation effect on the host.

We hypothesized that host MHC haplotypes would selectively target specific bacterial communities, co-evolving in a manner that increases host survival in the face of pathogenic infections. To this extent, no infection data exist to fully test this hypothesis and further investigations in this regard are needed. However, one of our main results of this study indicates that individuals carrying supertype_haplotype_ 1_3, 2_2, 2_4 and 3_3 have a specific bacterial composition. Besides, supertype 2 and 3, the most abundant supertypes in south and north, respectively, are antagonistically linked to specific bacterial taxa. For instance, taxonomic units ASV52, ASV73 and ASV71 that are included within the proteobacteria group that are from the Oxalobacteraceae family, are positively correlated with Supertype 3 but negatively correlated with Supertype 2. Bacteria from family Oxalobacteraceae have been recently detected in amphibian skin among individuals with different Bd infection intensity rates in amphibian (Ellison, Knapp, Sparagon, Swei, & Vredenburg, 2019). This result might indicate a different strategy to combat infectious diseases between regions. We suggest that specific bacteria from Oxalobacteraceae family could act differently on infected individuals depending of their MHC class II supertype configuration and bacterial abundance, but this deserves further investigation.

Together, these findings suggest that the evolutionary associations between host and microbiota is a complex evolutionary process modulated by distinct historical processes, local environmental conditions, and genetic characteristics of the host. Several studies support the idea that local environmental conditions might directly predict the amphibian skin microbiome structure by influencing the pool of potential symbionts in the habitat (Amato et al., 2016; Kueneman et al., 2014; Rebollar et al., 2016), but none of them considered the evolutionary history of populations, and how drift, local adaptation, and gene flow affect the host genome and how this affects microbiome composition. Therefore, our study shows that a combination of 1) evolutionary and biogeographic processes, 2) local environmental conditions and 3) host genome characteristics, may contribute to shape the skin microbiota diversity and heterogeneity. The study of these factors is essential for understanding host-microbiome-immunity interactions. Further surveying of wild populations along environmental gradients may help to identify environmental characteristics and evolutionary processes that shape host-associated microbial communities.

## Acknowledgements

We thank David Åhlen for the invaluable help in the field. Many thanks to Gunilla Egström for all the help and support in the lab. We also want to thank Javier Edo Varg for the help with microbiome pipeline analysis and all the guidance in the lab and to Yvonne Meyer-Lucht for the invaluable help during the study design.

## Data accessibility

Electronic supplementary material is available online: https://figshare.com/s/fa4e49bd4aa8e9f8819b

Raw data avaible from NCBI: pending

## Authors contribution

MC; conceptualization, field work, lab work, data curation, formal analyses, writing-original draft, funding; ARB.; field work, conceptualization, formal analyses, writing-original draft, review and editing, PRM; field work, statistical-bioinformatic support, writing – review and editing, PH; statistical-bioinformatic support, writing-review and editing; JBL: field work, writing-review and editing; AL; conceptualization, funding, writing-review and editing, JH; conceptualization, funding, writing-review and editing.

