## Supplementary material for "Association between the skin microbiome and MHC class II diversity in an amphibian": suplementary Material

**Suplementary material: Tables, Additional information and Figures**

**Table S1.** Information about pond locations within the two regions of the study: Uppsala (south) and Luleå (north)

| **Region** | **Population** | **Name** | **Ycoord** | **Xccord** |
| --- | --- | --- | --- | --- |
| Uppsala | p10 | Kroklösa | 59.617197 | 16.923972 |
| Uppsala | p1 | Ekeborg | 59.730955 | 16.987833 |
| Uppsala | p4 | Valsbrunna | 59.754256 | 17.037403 |
| Uppsala | p3 | Eneby | 59.741369 | 16.839627 |
| Uppsala | p22 | Högbyhatt | 59.820436 | 17.564300 |
| Uppsala | p23 | Dalkarlskärret | 59.776084 | 17.584391 |
| Uppsala | p18 | Mosta | 59.931544 | 17.297956 |
| Uppsala | p26 | Ströbykärret | 59.922572 | 17.307414 |
| Luleå | p34 | Lillträsket | 65.583369 | 22.318827 |
| Luleå | p35 | Vittjärnen | 65.520554 | 21.687510 |
| Luleå | p36 | Djurhustjärnen | 65.509020 | 21.717596 |
| Luleå | p37 | Dalbacka | 65.488997 | 21.378265 |

**Table S2. Illumina i7 and i5 indexes used in this study**

| **Illumina Index** |  | **i7 sequence** |
| --- | --- | --- |
| **705** | GGACTCCT | CAAGCAGAAGACGGCATACGAGATGGACTCCTGTGACTGGAGTTCAGACGTGTGCTCTTCCGATCT |
| **706** | TAGGCATG | CAAGCAGAAGACGGCATACGAGATTAGGCATGGTGACTGGAGTTCAGACGTGTGCTCTTCCGATCT |
| **707** | CTCTCTAC | CAAGCAGAAGACGGCATACGAGATCTCTCTACGTGACTGGAGTTCAGACGTGTGCTCTTCCGATCT |
| **708** | CAGAGAGG | CAAGCAGAAGACGGCATACGAGATCAGAGAGGGTGACTGGAGTTCAGACGTGTGCTCTTCCGATCT |
| **709** | GCTACGCT | CAAGCAGAAGACGGCATACGAGATGCTACGCTGTGACTGGAGTTCAGACGTGTGCTCTTCCGATCT |
| **710** | CGAGGCTG | CAAGCAGAAGACGGCATACGAGATCGAGGCTGGTGACTGGAGTTCAGACGTGTGCTCTTCCGATCT |
| **711** | AAGAGGCA | CAAGCAGAAGACGGCATACGAGATAAGAGGCAGTGACTGGAGTTCAGACGTGTGCTCTTCCGATCT |
| **712** | GTAGAGGA | CAAGCAGAAGACGGCATACGAGATGTAGAGGAGTGACTGGAGTTCAGACGTGTGCTCTTCCGATCT |
| **733** | CTTATACC | CAAGCAGAAGACGGCATACGAGATCTTATACCGTGACTGGAGTTCAGACGTGTGCTCTTCCGATCT |
| **735** | TCTAGCTG | CAAGCAGAAGACGGCATACGAGATTCTAGCTGGTGACTGGAGTTCAGACGTGTGCTCTTCCGATCT |
| **736** | CCATAGCA | CAAGCAGAAGACGGCATACGAGATCCATAGCAGTGACTGGAGTTCAGACGTGTGCTCTTCCGATCT |
| **738** | GGTATAGC | CAAGCAGAAGACGGCATACGAGATGGTATAGCGTGACTGGAGTTCAGACGTGTGCTCTTCCGATCT |
| **739** | GGTTATGC | CAAGCAGAAGACGGCATACGAGATGGTTATGCGTGACTGGAGTTCAGACGTGTGCTCTTCCGATCT |
| **740** | TAGGCAAG | CAAGCAGAAGACGGCATACGAGATTAGGCAAGGTGACTGGAGTTCAGACGTGTGCTCTTCCGATCT |
| **741** | TTGTCCAT | CAAGCAGAAGACGGCATACGAGATTTGTCCATGTGACTGGAGTTCAGACGTGTGCTCTTCCGATCT |
| **743** | TCTAGGCA | CAAGCAGAAGACGGCATACGAGATTCTAGGCAGTGACTGGAGTTCAGACGTGTGCTCTTCCGATCT |
| **701** | TCGCCTTA | CAAGCAGAAGACGGCATACGAGATTCGCCTTAGTGACTGGAGTTCAGACGTGTGCTCTTCCGATCT |
| **702** | CTAGTACG | CAAGCAGAAGACGGCATACGAGATCTAGTACGGTGACTGGAGTTCAGACGTGTGCTCTTCCGATCT |
| **703** | AGGCAGAA | CAAGCAGAAGACGGCATACGAGATAGGCAGAAGTGACTGGAGTTCAGACGTGTGCTCTTCCGATCT |
| **704** | TCCTGAGC | CAAGCAGAAGACGGCATACGAGATTCCTGAGCGTGACTGGAGTTCAGACGTGTGCTCTTCCGATCT |
| **N715** | CCTGAGAT | CAAGCAGAAGACGGCATACGAGATCCTGAGATGTGACTGGAGTTCAGACGTGTGCTCTTCCGATCT |
| **N716** | TAGCGAGT | CAAGCAGAAGACGGCATACGAGATTAGCGAGTGTGACTGGAGTTCAGACGTGTGCTCTTCCGATCT |
| **N718** | GTAGCTCC | CAAGCAGAAGACGGCATACGAGATGTAGCTCCGTGACTGGAGTTCAGACGTGTGCTCTTCCGATCT |
| **N719** | TACTACGC | CAAGCAGAAGACGGCATACGAGATTACTACGCGTGACTGGAGTTCAGACGTGTGCTCTTCCGATCT |
| **N720** | AGGCTCCG | CAAGCAGAAGACGGCATACGAGATAGGCTCCGGTGACTGGAGTTCAGACGTGTGCTCTTCCGATCT |
| **N721** | GCAGCGTA | CAAGCAGAAGACGGCATACGAGATGCAGCGTAGTGACTGGAGTTCAGACGTGTGCTCTTCCGATCT |
| **N722** | CTGCGCAT | CAAGCAGAAGACGGCATACGAGATCTGCGCATGTGACTGGAGTTCAGACGTGTGCTCTTCCGATCT |
| **N723** | GAGCGCTA | CAAGCAGAAGACGGCATACGAGATGAGCGCTAGTGACTGGAGTTCAGACGTGTGCTCTTCCGATCT |
|  |  | **5i sequence** |
| **521** | CTTGCTTT | AATGATACGGCGACCACCGAGATCTACACCTTGCTTTACACTCTTTCCCTACACGACG |
| **522** | GGCTTCAA | AATGATACGGCGACCACCGAGATCTACACGGCTTCAAACACTCTTTCCCTACACGACG |
| **523** | AATCGGCA | AATGATACGGCGACCACCGAGATCTACACAATCGGCAACACTCTTTCCCTACACGACG |
| **524** | GGTTCAAA | AATGATACGGCGACCACCGAGATCTACACGGTTCAAAACACTCTTTCCCTACACGACG |
| **525** | ACTTCGAC | AATGATACGGCGACCACCGAGATCTACACACTTCGACACACTCTTTCCCTACACGACG |
| **526** | TGACTTGC | AATGATACGGCGACCACCGAGATCTACACTGACTTGCACACTCTTTCCCTACACGACG |
| **527** | TAGGACCT | AATGATACGGCGACCACCGAGATCTACACTAGGACCTACACTCTTTCCCTACACGACG |
| **528** | GGAGACTT | AATGATACGGCGACCACCGAGATCTACACGGAGACTTACACTCTTTCCCTACACGACG |

**TableS3**. Descriptive table of MHC and Skin microbiome diversity indexes in the two regions (Uppland and Luleå) included in the study. N= Number of individuals collected per population, N_heterozygous_ = Number of heterozygous individuals, N_homozygous_ = Number of homozygous individuals, AR_h_=Allelic Richness per haplotype, AR_S_=allelic richness per supertype, HO= observed homozygosity, HE=expected homozygosity, π = nucleotide diversity, theta K = average number of nucleotide differences, Alpha diversity (Shannon, Simpson and Chao1).

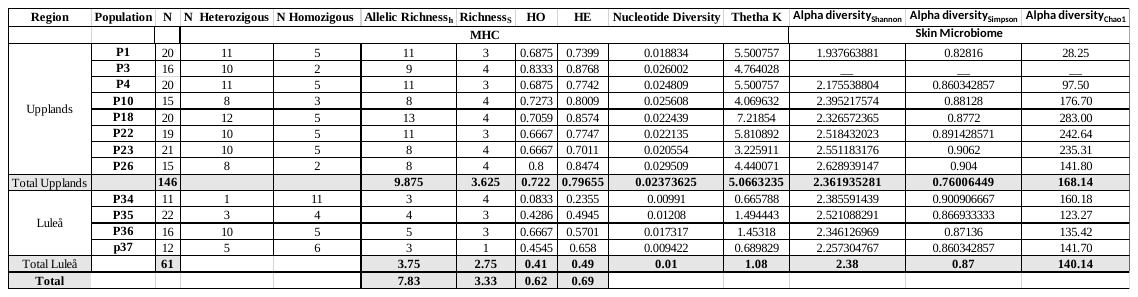

**Table S4.** PERMANOVA test showing significant differences in bacterial community composition between two different types of samples: water filters and adult swabs. adonis2 () function of vegan package was implemented. A Significant p-value is highlighted with (*)

|  | **Df** | **SumsOfSqs** | **MeanSqs** | **F.Model** | **R2** | **Pr(>F)** |  |
| --- | --- | --- | --- | --- | --- | --- | --- |
| **~sample_type** | 1 | 21922 | 21922 | 3.8262 | 0.02176 | 0.001 | *** |
| **Residuals** | 172 | 985465 | 5729.4 |  | 0.97824 |  |  |
| **Total** | 173 | 1007387 |  |  | 1 |  |  |

**Table S5.** Simple General linear models (GLM) showing that temperature at data collections (tempCollection) has an effect on skin microbiome diversity (Shannon diversity index) as well as Mean average precipitation (PreMean). Generalized linear models (GLMM) show that the Mean average precipitation (PreMean) and for Region were positively related to alpha-diversity (Shannon diversity index). lmer () function of lme4 package was implemented. Significant p-value is highlighted with (*).

| **GLM; aov** | **Df** | **Sum Sq** | **Mean Sq** | **F value** | **Pr(>F)** |  |
| --- | --- | --- | --- | --- | --- | --- |
| Region | 1.00 | 0.01 | 0.01 | 0.01 | 0.01 |  |
| temCollection | 1.00 | 2.83 | 2.83 | 3.76 | *0.05* | . |
| PreMean | 1.00 | 9.35 | 9.35 | 12.41 | **0.00** | *** |
| region*temCollection | 1.00 | 5.19 | 5.19 | 6.88 | **0.01** | ** |
| Region*PreMean | 1.00 | 0.51 | 0.51 | 0.68 | 0.41 |  |
| Residuals | 158.00 | 118.99 | 0.75 |  |  |  |

| **GLMM; lmer** |  |  |  |  |  |  |  |
| --- | --- | --- | --- | --- | --- | --- | --- |
| **Random effects:** |  |  |  |  |  |  |  |
| Groups | Name | Variance | Std.Dev. |  |  |  |  |
| PopNAME | (Intercept) | 0.09 | 0.3 |  |  |  |  |
| Residual |  | 0.7 | 0.83 |  |  |  |  |
| Number of obs: | 164 |  |  |  |  |  |  |
| groups:PopNAME | 12 |  |  |  |  |  |  |
| **Fixed effects:** | Estimate | Std. Error | df | t value | Pr(>\|t\|) |  |  |
| (Intercept) | 38.05 | 15.35 | 6.34 | 2.48 | **0.046** | * |  |
| Region_south | 7.77 | 3.47 | 7.26 | 2.24 | 0.05 | . |  |
| PreMean | -0.51 | 0.21 | 6.07 | -2.44 | **0.049** | * |  |
| TemCollection | 1.89 | 2.34 | 6.66 | 0.81 | 0.44 |  |  |
| TemMean | -3.61 | 3.01 | 6.75 | -1.2 | 0.26 |  |  |
| **Correlation of Fixed Effects:** | (Intr) | rgnsth | PreMen | tmCllc |  |  |  |
| Region_south | 0.88 |  |  |  |  |  |  |
| PreMean | -0.95 | -0.78 |  |  |  |  |  |
| TemCollection | -0.85 | 0.41 | -0.65 |  |  |  |  |
| temMean | -0.85 | -0.62 | 0.76 | -0.96 |  |  |  |
| **Type III Analysis of Variance Table with Satterthwaite's method** | SumSq | Mean Sq | NumbeDF | DenDF | F value | Pr(>F) |  |
| Region | 3.51 | 3.51 | 1 | 7.26 | 5.02 | 0.058 | . |
| PreMean | 4.17 | 4.17 | 1 | 6.07 | 5.96 | **0.049** | * |
| TemCollection | 0.45 | 0.45 | 1 | 6.66 | 0.65 | 0.44 |  |
| TemMean | 1.01 | 1.01 | 1 | 6.75 | 1.44 | 0.26 |  |

**Table S6.** PERMANOVA test showing significant differences in bacterial community composition between the two regions, by using *adonis2()* function of vegan package. Significant p-values are highlighted in bold (* p-value between 0.05 and 0.01, ** p-value between 0.01 and 0.0001, *** <0.0001).

|  | **Df** | **SumsOfSqs** | **MeanSqs** | **F.Model** | **R^2^** | **Pr(>F)** |
| --- | --- | --- | --- | --- | --- | --- |
| **Adonis() ~region** | 1 | 27152 | 27152.2 | 4.8705 | 0.02919 | **0.001***** |
| **Residuals** | 162 | 903115 | 5574.8 |  | 0.97081 |  |
| **Total** | 163 | 930267 |  |  | 1 |  |

**Table S7.** Permutation test (PERMADIST) for homogeneity of multivariate dispersions.

|  | **Df** | **Sum Sq** | **Mean Sq** | **F** | **N.Perm** | **Pr(>F)** |
| --- | --- | --- | --- | --- | --- | --- |
| **Groups** | 1 | 1673 | 1672.8 | 1.9223 | 999 | 0.169 |
| **Residuals** | 162 | 140975 | 870.21 |  |  |  |

**Table S8. Results from** Multiple regression (MRM) and linear regression models (lm) of relationship between Shannon diversity index and MHC heterozygosity/Nucleotide diversity Statistical models included Shannon diversity index as the response variable and diversity index as a fixed factor. Significant p-values are highlighted in bold (** p-value <0.01 and *** p-value <0.0001).

| **MRM multiple regressions** | **Shannon diversity index** | **p-value** |  | **R^2^** | **p-value** | **F.test** | **p-value** |  |  |
| --- | --- | --- | --- | --- | --- | --- | --- | --- | --- |
| Intercept | 0.23 | **0.000** | ******* | 0.34 | **0.0001** | 27.59 | **<0.001** | ******* |  |
| MHCHeterozygosity | 4.52 | **0.000** | ******* |  |  |  |  |  |  |
| Intercept | **Shannon diversity index** | **p-value** |  | **R^2^** | **p-value** | **F.test** | **p-value** |  |  |
| MHCHeterozygosity | 0.28 | 0.99 |  | 0.355 | **0.0002** | 14.32 | **<0.001** | ******* |  |
| Geographical distance | 4.46 | **0.0001** | ******* |  |  |  |  |  |  |
|  | -0.0001 | 0.31 |  |  |  |  |  |  |  |
| Intercept | **Shannon diversity index** | **p-value** |  | **R^2^** | **p-value** | **F.test** | **p-value** |  |  |
| MHC Nucleotide diversity | 5.83 | **0.0036** | ******* | 0.0008 | **0.0096** | 9.65 | **0.010** | ****** |  |
|  | -0.64 | **0.0096** | ******* |  |  |  |  |  |  |
| **Linear Regression (lm)** | **Estimate** | **Std. Error** |  | **t value** | **Pr(>\|t\|)** | **Fvalue** | **DF** | **p-value** |  |
| Intercept | 0.23 | **0.85** |  | 2.71 | **0.0091** |  |  |  |  |
| MHCHeterozygosity | 4.52 | 0.86 |  | 5.25 | **2.7E-06** | 27.3 | 53 | **<0.001** | ******* |
| Intercept | 5.83 | 0.05585 |  | 104.46 | **<2e-16** |  |  |  |  |
| Nucleotide diversity | -0.64 | 0.20874 |  | -3.11 | **0.0019** | 9.65 | 11323 | **<0.001** | ******* |

**Table S9.** A General Lineal Model (GLM) using the car package in R showing the *Shannon diversity* index was the response variable and the *region* the fix factor. Data was subset by MHC class II exon 2 *Heterozygosity* corresponding to only individuals that belong to the *DistinctS* group. P-value <0.05 is indicated by *.

| **Response: Shannon diversity index** | **Df** | **Sum Sq** | **Mean sq** | **Fvalue** | **Pr(>F)** |  |
| --- | --- | --- | --- | --- | --- | --- |
| **Region** | 1 | 0.8839 | 0.88393 | 4.1986 | **0.04524** | ***** |
| **Residuals** | 55 | 11.5791 | 0.21053 |  |  |  |

**Table S10.** dbRDA multivariate statistics model showing significant differences in skin microbiome composition between supertype_haplotypes. Significant p-values are highlighted in bold (** p-value between 0.01 and 0.0001).

|  | **Df** | **Inertia** | **F** | **Pr(>F)** |
| --- | --- | --- | --- | --- |
| **Model** | 1 | 0.4653 | 6.1771 | **0.00222 **** |
| **Residual** | 146 | 10.9985 |  |  |

**Table S11.** PERMANOVA; *Adonis2*() test showing that individuals carrying supertype H1 and H2 present a different skin bacterial composition. Significant supertypes p-values are highlighted in bold (* p-value between 0.05 and 0.01, ** p-value between 0.01 and 0.0001, *** <0.0001).

|  | **Df** | **SumOfSqs** | **R2** | **F** | **Pr(>F)** |  |
| --- | --- | --- | --- | --- | --- | --- |
| **S1** | 1 | 0.889 | 0.01485 | 2.352 | **0.00444** | ****** |
| **S2** | 1 | 1.422 | 0.02374 | 3.7611 | **0.00111** | ****** |
| **S3** | 1 | 0.459 | 0.00766 | 1.2137 | 0.20089 |  |
| **S4** | 1 | 0.408 | 0.00681 | 1.0792 | 0.30855 |  |
| **Residual** | 150 | 56.721 | 0.94693 |  |  |  |
| **Total** | 154 | 59.9 | 1 |  |  |  |

**Table S12.** PERMANOVA; *Adonis2*() test show significant differences in skin microbiome composition depending on the specific supertype_haplotype. Significant supertype_haplotypes are marked with an (*).

|  | **Df** | **SumOfSqs** | **R2** | **F** | **Pr(>F)** |  |
| --- | --- | --- | --- | --- | --- | --- |
| S1_1 | 1 | 0.498 | 0.00832 | 1.3257 | 0.139845 |  |
| S1_2 | 1 | 0.377 | 0.00629 | 1.0021 | 0.437292 |  |
| S1_3 | 1 | 0.728 | 0.01215 | 1.9374 | 0.013319 | * |
| S2_2 | 1 | 1.021 | 0.01705 | 2.7172 | 0.00333 | ** |
| S2_3 | 1 | 0.369 | 0.00616 | 0.9821 | 0.431743 |  |
| S2_4 | 1 | 0.75 | 0.01252 | 1.9965 | 0.006659 | ** |
| S3_3 | 1 | 0.841 | 0.01404 | 2.2379 | 0.006659 | ** |
| S3_4 | 1 | 0.455 | 0.00759 | 1.21 | 0.208657 |  |
| Residual | 146 | 54.861 | 0.91588 |  |  |  |
| Total | 154 | 59.9 | 1 |  |  |  |

**Table S13.** ASVs differentially abundant per supertype by DESEq2 software. ASVs that coincided between different approaches (DESEq2 and ANCOMBC2) between supertypes, are highlighted in and grey (supertype2) and green (supertype3), respectively. This ASVs represent the taxa grouped by *Genus* that are shown significantly different in abundance when the ANCOMBC software is used.

| **DESEq2** | | |
| --- | --- | --- |
| **ASVs ID** | **Taxa information** | **padj** |
| ASV50 | P; Bacteroidetes, C; Flavobacteriia, O; Flavobacteriale, F; Flavobacteriaceae, G; Chryseobacterium | 8.87E-20 |
| ASV48 | P; Bacteroidetes, C; Flavobacteriia, O; Flavobacteriale, F; Flavobacteriaceae, G; Chryseobacterium | 4.37E-18 |
| ASV62 | P Proteobacteria, C; Gammaproteobacteria, O; Pseudomonadales, F; Pseudomonadaceae, Pseudomonas | 5.62E-16 |
| **supertype 2 (ASV)** | | |
| ASV62 | P;Proteobacteria, C; Gammaproteobacteria, O; Pseudomonadales, F; Pseudomonadaceae, G; Pseudomonas | 1.09E-50 |
| ASV80 | P;Proteobacteria, C; Gammaproteobacteria, O; Pseudomonadales, F; Pseudomonadaceae, G; Pseudomonas | 4.95E-46 |
| ASV57 | P;Proteobacteria, C; Gammaproteobacteria, O; Pseudomonadales, F; Pseudomonadaceae, G; Pseudomonas | 1.61E-45 |
| ASV72 | P;Proteobacteria, C; Gammaproteobacteria, O; Pseudomonadales, F; Pseudomonadaceae, G; Pseudomonas | 4.77E-39 |
| ASV77 | P;Proteobacteria, C; Betaproteobacteria, O; Burkholderiales; F; Comamonadaceae, G; Polaromonas | 4.96E-35 |
| ASV89 | P;Proteobacteria, C; Betaproteobacteria, O; Burkholderiales; F; Comamonadaceae, G; Polaromonas | 4.96E-35 |
| ASV95 | P;Proteobacteria, C; Betaproteobacteria, O; Burkholderiales; F; Comamonadaceae, G; Polaromonas | 4.00E-30 |
| ASV63 | P;Proteobacteria, C; Betaproteobacteria, O; Burkholderiales; F; Comamonadaceae; G; Rhodoferax | 9.09E-03 |
| ASV76 | P;Proteobacteria, C; Betaproteobacteria, O; Burkholderiales; F; Oxalobacteraceae, G; Herbaspirillum | 1.35E-02 |
| ASV65 | P;Proteobacteria, C; Betaproteobacteria, O; Burkholderiales; F; Comamonadaceae; G; Rhodoferax | 1.55E-02 |
| ASV216 | P;Proteobacteria, C; Betaproteobacteria, O; Burkholderiales; F; Oxalobacteraceae, G; Herbaspirillum | 3.19E-02 |
| ASV36 | P;Proteobacteria, C; Betaproteobacteria, O; Burkholderiales, F;Comamonadaceae, G; Rhodoferax | 3.19E-02 |
| ASV53 | P;Proteobacteria, C; Betaproteobacteria, O; Burkholderiales; F; Oxalobacteraceae, G; Herbaspirillum | 3.19E-02 |
| ASV56 | P;Proteobacteria, C; Betaproteobacteria, O; Burkholderiales; F; Oxalobacteraceae, G; Herbaspirillum | 3.19E-02 |
| ASV79 | P;Proteobacteria, C; Betaproteobacteria, O; Burkholderiales; F; Oxalobacteraceae, G; Herbaspirillum | 3.19E-02 |
| ASV86 | P;Proteobacteria, C; Betaproteobacteria, O; Burkholderiales, F; Burkholderiales incertae sedis, G; Mitsuaria | 3.19E-02 |
| ASV51 | P;Proteobacteria, C; Betaproteobacteria, O; Burkholderiales; F; Oxalobacteraceae, G; Undibacterium | 3.98E-02 |
| ASV59 | P;Proteobacteria, C; Betaproteobacteria, O; Burkholderiales, F; Burkholderiales incertae sedis, G; Mitsuaria | 4.35E-02 |
| ASV83 | P;Proteobacteria; C; Betaproteobacteria, O; Burkholderiales; F; Oxalobacteraceae, G;Herminiimonas | 4.80E-02 |
| ASV64 | P;Proteobacteria, C; Betaproteobacteria, O; Burkholderiales, F; Burkholderiales incertae sedis, G; Mitsuaria | 4.87E-02 |
| **Supertype3 (ASV)** | | |
| ASV62 | P;Proteobacteria_C;Gammaproteobacteria_O;Pseudomonadales_F;Pseudomonadaceae_Pseudomonas | 9.97E-52 |
| ASV80 | P;Proteobacteria_C;Gammaproteobacteria_O;Pseudomonadales_F;Pseudomonadaceae_Pseudomonas | 6.02E-47 |
| ASV72 | P;Proteobacteria_C;Gammaproteobacteria_O;Pseudomonadales_F;Pseudomonadaceae_Pseudomonas | 7.57E-47 |
| ASV57 | P;Proteobacteria_C;Gammaproteobacteria_O;Pseudomonadales_F;Pseudomonadaceae_Pseudomonas | 9.66E-47 |
| ASV89 | P;Proteobacteria_C;Betaproteobacteria_O;Burkholderiales_F;Comamonadaceae_Polaromonas | 2.08E-36 |
| ASV77 | P;Proteobacteria_C;Betaproteobacteria_O;Burkholderiales_F;Comamonadaceae_Polaromonas | 1.30E-31 |
| ASV52 | P;Proteobacteria_C;Betaproteobacteria_O;Burkholderiales_F;Oxalobacteraceae_Undibacterium | 9.45E-06 |
| ASV71 | P;Proteobacteria_C;Betaproteobacteria_O;Burkholderiales_F;Oxalobacteraceae_Undibacterium | 2.46E-05 |
| ASV73 | P;Proteobacteria_C;Betaproteobacteria_O;Burkholderiales_F;Oxalobacteraceae_Undibacterium | 2.46E-05 |
| ASV51 | P;Proteobacteria_C;Betaproteobacteria_O;Burkholderiales_F;Oxalobacteraceae_Undibacterium | 7.84E-05 |
| ASV33 | P;Bacteroidetes_C;Flavobacteriia_O;Flavobacteriales_F;Flavobacteriaceae_Chryseobacterium | 5.26E-04 |
| ASV50 | P;Bacteroidetes_C;Flavobacteriia_O;Flavobacteriales_F;Flavobacteriaceae_Chryseobacterium | 8.03E-04 |
| ASV48 | P;Bacteroidetes_C;Flavobacteriia_O;Flavobacteriales_F;Flavobacteriaceae_Chryseobacterium | 1.19E-03 |
| ASV37 | P;Proteobacteria_C;Betaproteobacteria_O;Burkholderiales_F;Oxalobacteraceae_Herminiimonas | 1.21E-03 |
| ASV63 | P;Proteobacteria_C;Betaproteobacteria_O;Burkholderiales_F;Comamonadaceae_Rhodoferax | 1.47E-03 |
| ASV2 | P;Proteobacteria_C;Gammaproteobacteria_O;Pseudomonadales_F;Pseudomonadaceae_Pseudomonas | 2.24E-03 |
| ASV21 | P;Proteobacteria_C;Betaproteobacteria_O;Burkholderiales_F;Oxalobacteraceae_Herminiimonas | 4.39E-03 |
| ASV15 | P;Proteobacteria_C;Betaproteobacteria_O;Burkholderiales_F;Oxalobacteraceae_Herminiimonas | 5.01E-03 |
| ASV32 | P;Bacteroidetes_C;Flavobacteriia_O;Flavobacteriales_F;Flavobacteriaceae_Chryseobacterium | 7.05E-03 |
| ASV35 | P;Proteobacteria_C;Betaproteobacteria_O;Burkholderiales_F;Oxalobacteraceae_Herminiimonas | 8.83E-03 |
| ASV292 | P;Proteobacteria_C;Betaproteobacteria_O;Burkholderiales_F;Burkholderiales incertae sedis_Sphaerotilus | 1.69E-02 |
| ASV36 | P;Proteobacteria_C;Betaproteobacteria_O;Burkholderiales_F;Comamonadaceae_Rhodoferax | 1.72E-02 |
| ASV64 | P;Proteobacteria_C;Betaproteobacteria_O;Burkholderiales_F;Burkholderiales incertae sedis_Mitsuaria | 1.95E-02 |
| ASV59 | P;Proteobacteria_C;Betaproteobacteria_O;Burkholderiales_F;Burkholderiales incertae sedis_Mitsuaria | 2.03E-02 |
| ASV86 | P;Proteobacteria_C;Betaproteobacteria_O;Burkholderiales_F;Burkholderiales incertae sedis_Mitsuaria | 2.03E-02 |
| ASV65 | P;Proteobacteria_C;Betaproteobacteria_O;Burkholderiales_F;Comamonadaceae_Rhodoferax | 2.03E-02 |
| ASV1 | P;Proteobacteria_C;Gammaproteobacteria_O;Pseudomonadales_F;Pseudomonadaceae_Pseudomonas | 2.11E-02 |
| ASV87 | P;Proteobacteria_C;Betaproteobacteria_O;Burkholderiales_F;Burkholderiales incertae sedis_Mitsuaria | 2.93E-02 |
| ASV417 | P;Bacteroidetes_C;Sphingobacteriia_O;Sphingobacteriales_F;Sphingobacteriaceae_Pedobacter | 4.10E-02 |
| **Supertype4 (ASV)** | | |
| ASV91 | P;Proteobacteria_C;Betaproteobacteria_O;Burkholderiales_F;Oxalobacteraceae_G;Herminiimonas | 5.61E-29 |
| ASV65 | P;Proteobacteria_C;Betaproteobacteria_O;Burkholderiales_F;Comamonadaceae_G;Rhodoferax | 1.01E-22 |
| ASV98 | P;Proteobacteria_C;Betaproteobacteria_O;Burkholderiales_F;Oxalobacteraceae_G;Herminiimonas | 4.12E-22 |
| ASV63 | P;Proteobacteria_C;Betaproteobacteria_O;Burkholderiales_F;Comamonadaceae_G;Rhodoferax | 3.48E-21 |
| ASV21 | P;Proteobacteria_C;Betaproteobacteria_O;Burkholderiales_F;Oxalobacteraceae_G;Herminiimonas | 5.81E-05 |
| ASV15 | P;Proteobacteria_C;Betaproteobacteria_O;Burkholderiales_F;Oxalobacteraceae_G;Herminiimonas | 8.35E-05 |
| ASV51 | P;Proteobacteria_C;Betaproteobacteria_O;Burkholderiales_F;Oxalobacteraceae_G;Undibacterium | 0.002709 |
| ASV37 | P;Proteobacteria_C;Betaproteobacteria_O;Burkholderiales_F;Oxalobacteraceae_G;Herminiimonas | 0.009723 |
| ASV71 | P;Proteobacteria_C;Betaproteobacteria_O;Burkholderiales_F;Oxalobacteraceae_G;Undibacterium | 0.041024 |
| ASV73 | P;Proteobacteria_C;Betaproteobacteria_O;Burkholderiales_F;Oxalobacteraceae_G;Undibacterium | 0.041024 |
| ASV35 | P;Proteobacteria_C;Betaproteobacteria_O;Burkholderiales_F;Oxalobacteraceae_G;Herminiimonas | 0.043525 |

**Table S14.** Table representing differentially abundant taxa grouped by *Genus* per supertype by ANCOMBC2 software. Highlighted files in grey and green represent bacterial grouped by *Genus* that are significantly different in abundance by using the DESEq2 software.

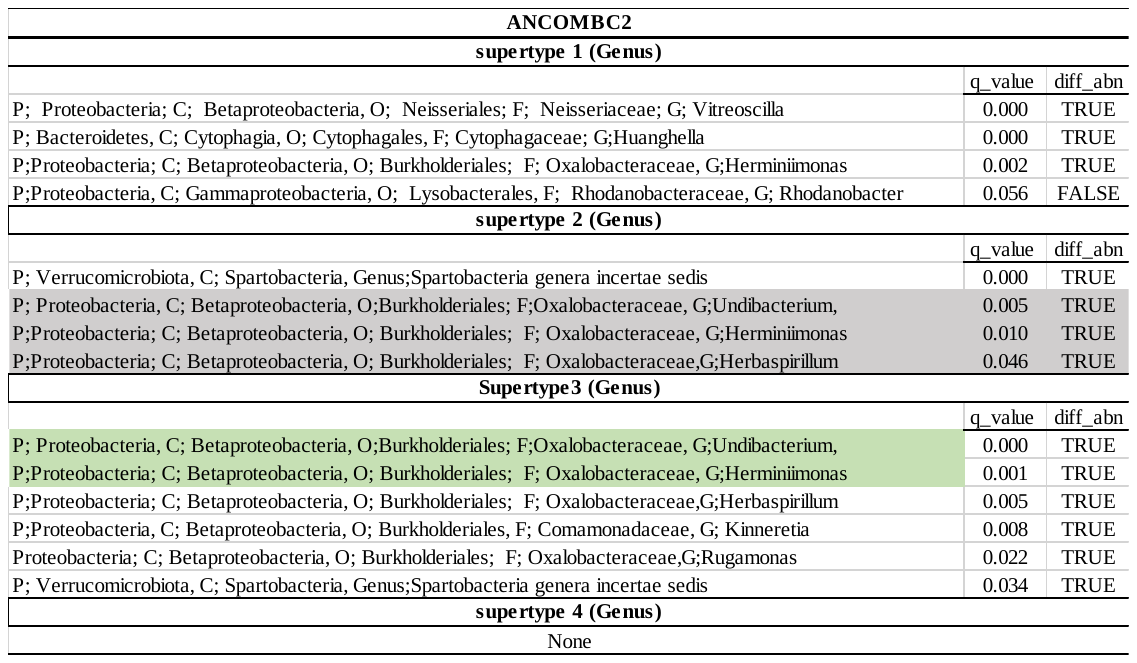

**Additional information A1**

**Skin microbiota PCR**

PCR reactions were performed in a 20-μl reaction volume comprising 1 μl of genomic DNA, 4 μl of 10X Phusion HF buffer (Thermo scientific lab), 0.4 μl of 2 mM of each dNTP, 0.25 μl of each 10 μM primer (515F and 806R, respectively), 0.6 μl of Bovine serum albumine (BSA; 5 mg/ml) and 0.2 μl of Phusion taq polymerase (5 U/μl, Thermo scientific lab) in deionized water. Thermocycling (GeneMax; Bioer Technology, Hangzhou, China) was conducted with an initial denaturation step at 98 °C for 30 s, followed by 30 cycles of denaturation at 98 °C for 10 s, annealing at 56 °C for 20 s and extension at 72 °C for 15 s, and finalised with a 8-min extension step at 72 °C. Triplicate technical replicates were run per sample, pooled after PCR amplification, and purified using the Agencourt AMPure XP purification kit (Beckman Coulter Inc., Brea, USA). In a second PCR-step, Illumina adaptors and multiplex identifiers (MIDs) were attached according to the final configuration. MIDs were 8 nucleotide long, sample specific, and developed following recommendations by Engelbrektson et al. (2010). PCR reactions were performed in a 30-uL reaction volume each reaction contained 0.5 μl of genomic DNA, 3 μl of 10X Phusion HF buffer), 0.6 μl of 2 mM of each dNTP, 0.75 μl of each 10 μM primer (515F and 806R, respectively), 0.6 μl of Bovine serum albumine (BSA; 5 mg/ml) and 0.3 μl of Phusion taq polymerase (5 U/μl, Thermo scientific lab) in deionized water. Each reaction started with initial denaturation at 98 °C for 30 s, followed by 20 cycles of denaturation at 98 °C for 10 s, annealing at 65 °C for 20 s and extension at 72 °C for 15 s, and finalized with a 8-min extension step at 72 °C. Again, triplicate technical replicates per sample were pooled, purified with the Agencourt AMPure XP purification kit (Beckman Coulter), and quantified on a fluorescence microplate reader (Ultra 384; Tecan Group Ltd., Männedorf, Switzerland), employing the Quant-iT PicoGreen dsDNA quantification kit (Invitrogen). Finally, PCR amplicons were pooled in equimolar proportions to obtain a similar number of sequencing reads per sample. The final, pooled amplicon was sequenced on a Illumina MiSeq system at the National Genomics Infrastructure by ScilifeLab, Uppsala, in Sweden.

Engelbrektson, A., Kunin, V., Wrighton, K. C., Zvenigorodsky, N., Chen, F., Ochman, H., & Hugenholtz, P. (2010). Experimental factors affecting PCR-based estimates of microbial species richness and evenness. The ISME journal, 4(5), 642-647.

**Figure S1**. The figure illustrates a positive relationship between the number of reads and observed taxonomic richness. Pearson correlation coefficients were used to estimate the strength of this correlation by using ggplot2 in R.

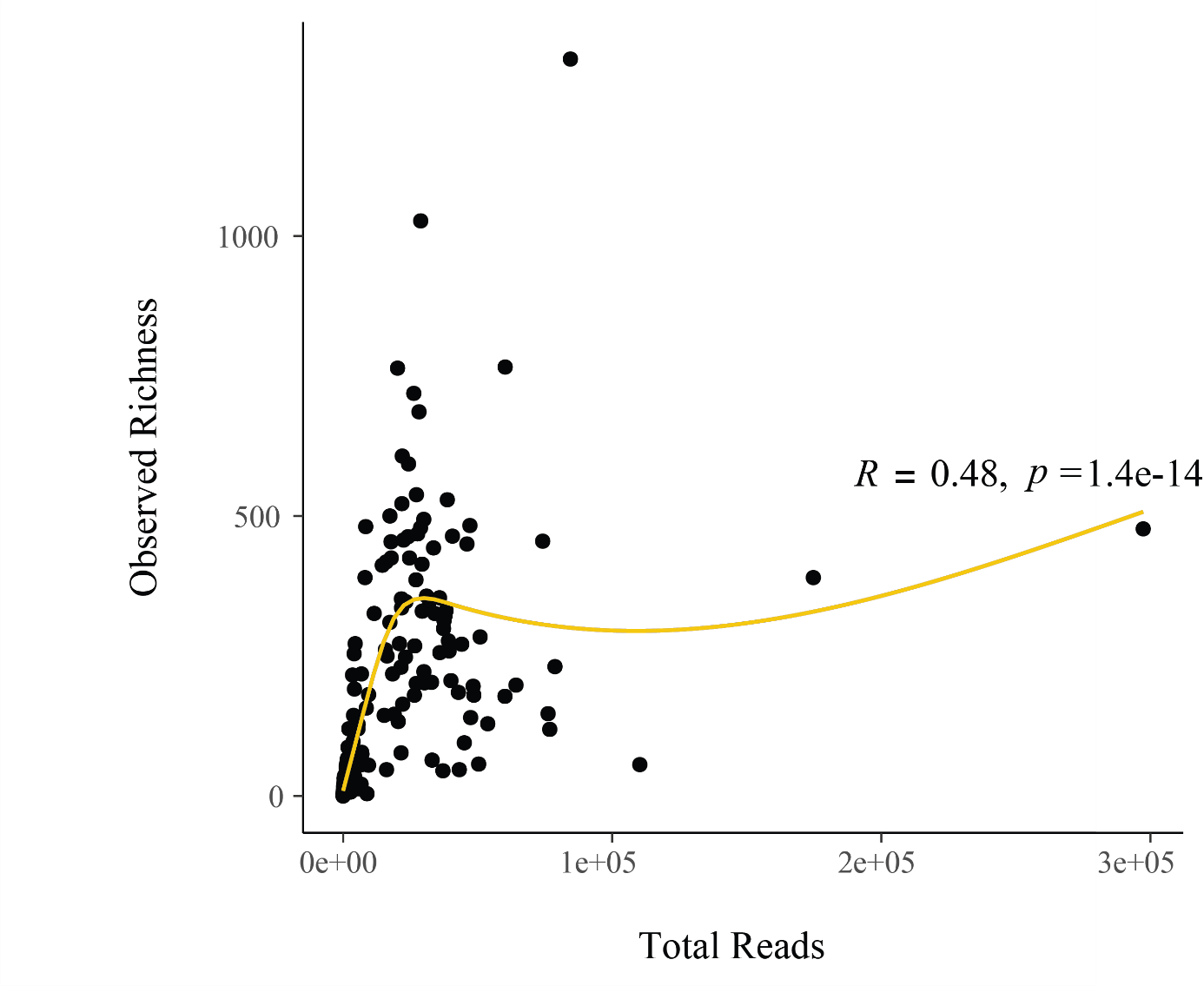

**Figure S2**. The abundance of skin bacteria at the phylum level based on the total number of sequences in our data set. A) for the amphibian samples and B) for the water filters.

A)

Amphibian samples

|  |  |
| --- | --- |
| Proteobacteria | 48.4003 |
| Bacteroidetes | 17.1401 |
| Actinobacteria | 10.4473 |
| Acidobacteria | 7.14985 |
| Verrucomicrobia | 5.69703 |
| Firmicutes | 3.91773 |
| Planctomycetes | 2.22005 |
| Chloroflexi | 1.87724 |
| Armatimonadetes | 1.48547 |
| Candidatus Saccharibacteria | 0.865165 |
| Gemmatimonadetes | 0.799869 |

B)

Water filters

|  |  |
| --- | --- |
| Proteobacteria | 36.5172 |
| Parcubacteria | 24.9272 |
| Candidatus Saccharibacteria | 16.8317 |
| Actinobacteria | 9.26034 |
| Chlamydiae | 3.20326 |
| Verrucomicrobia | 2.91206 |
| Firmicutes | 2.56261 |
| Acidobacteria | 2.38789 |
| Bacteroidetes | 1.39779 |

**Figure S3.** Color scheme for the allele distribution maps for the haplotypes and supertypes_haplotypes used in Figure 2.

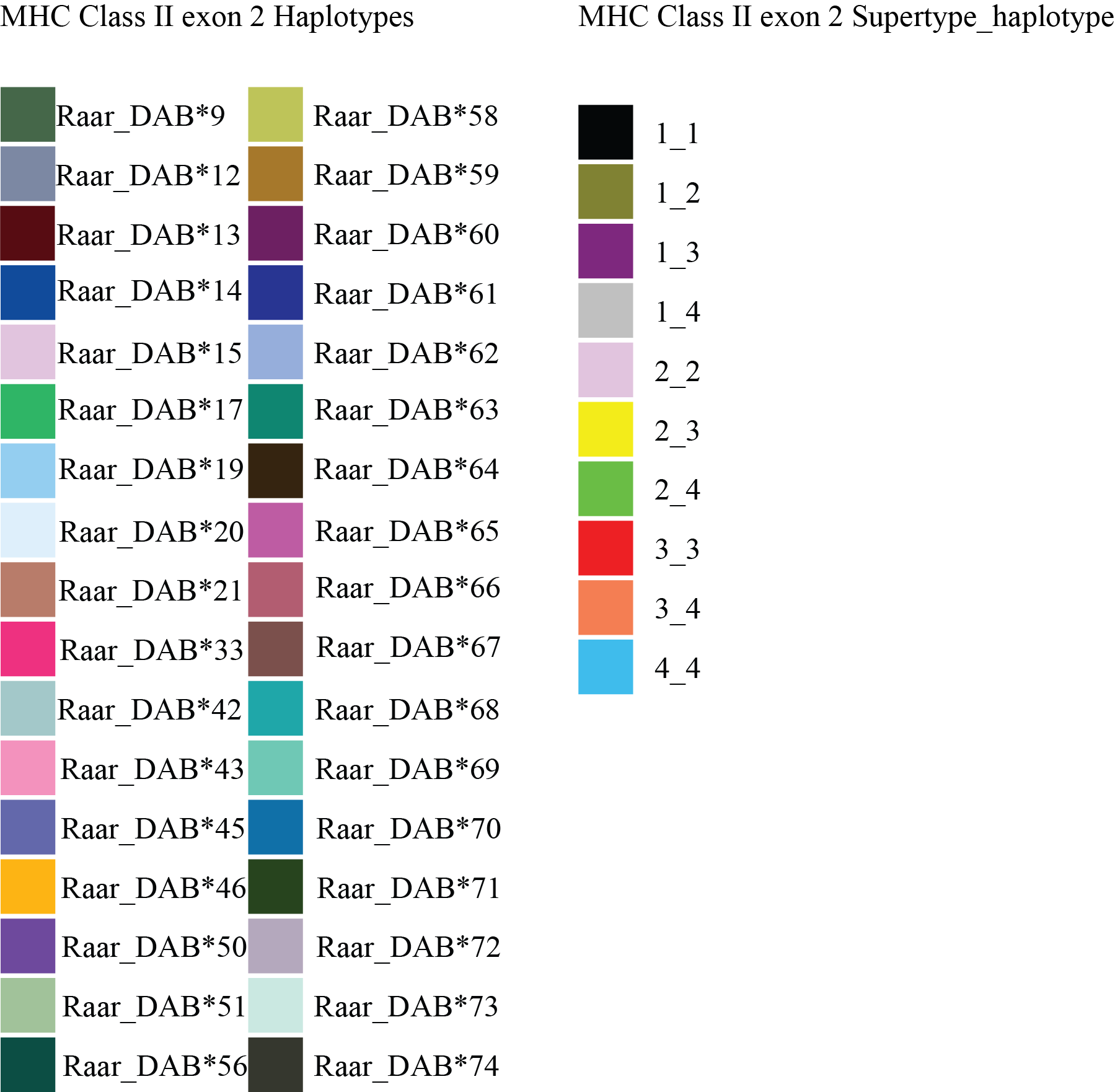

**Figure S4**. A) PCoA plot demonstrated the beta diversity of water filter microbial community and Adult swab samples at ASVs level on weighted UniFrac distance derived from 16S microbial sequences (PERMANOVA; Adonis2, P <0.05). B) Represent the distance to the polygon centroids (PERMADIST; p<0.05).

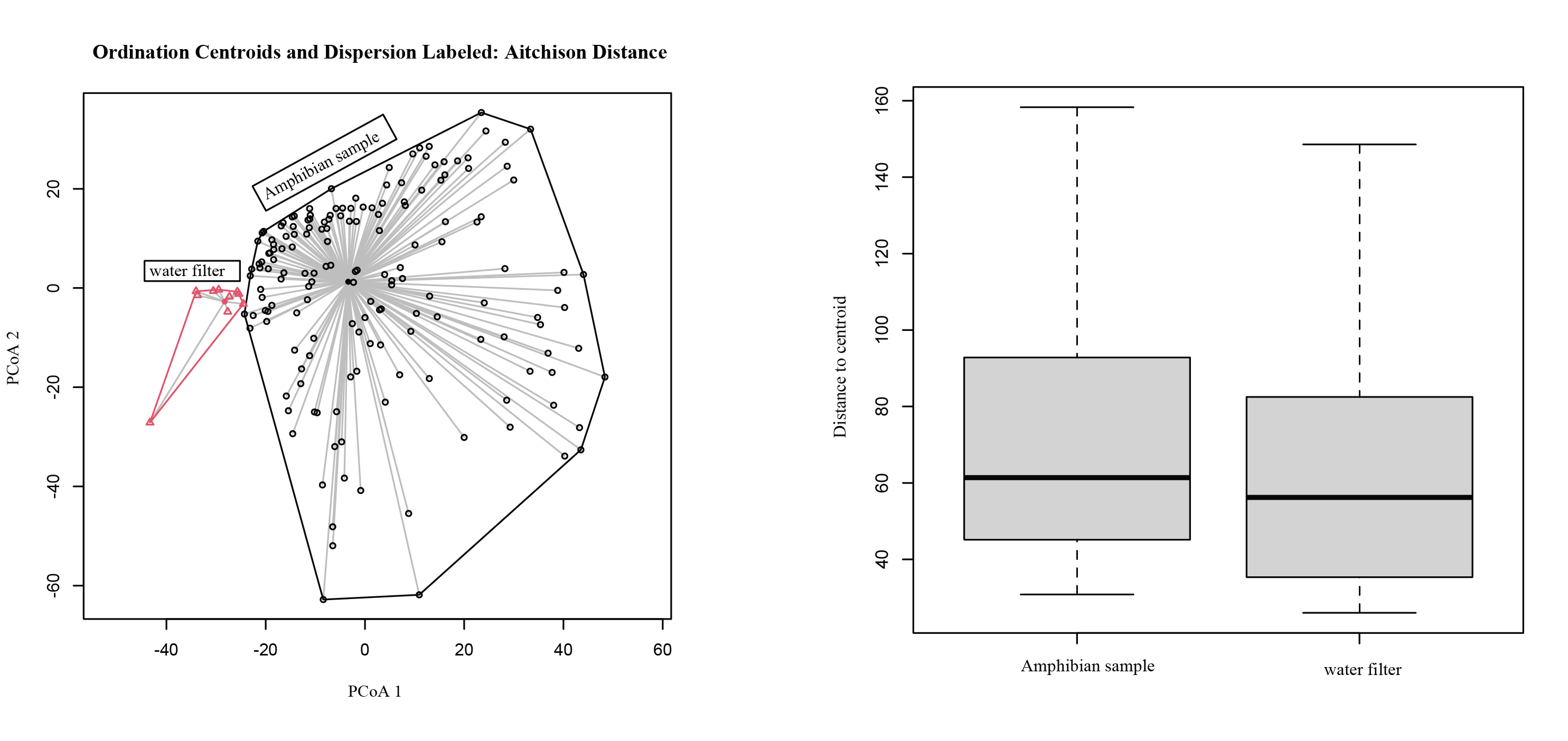

**A**

**B**

**Figure S5**. Alpha Diversity index (Observed, Shannon and PD) at the two sampling sites (South) Uppland and (North) Luleå. The purple circles represent all the 16s skin microbial composition samples from Uppland while the orange circles represent the 16s skin microbial composition samples from Luleå.

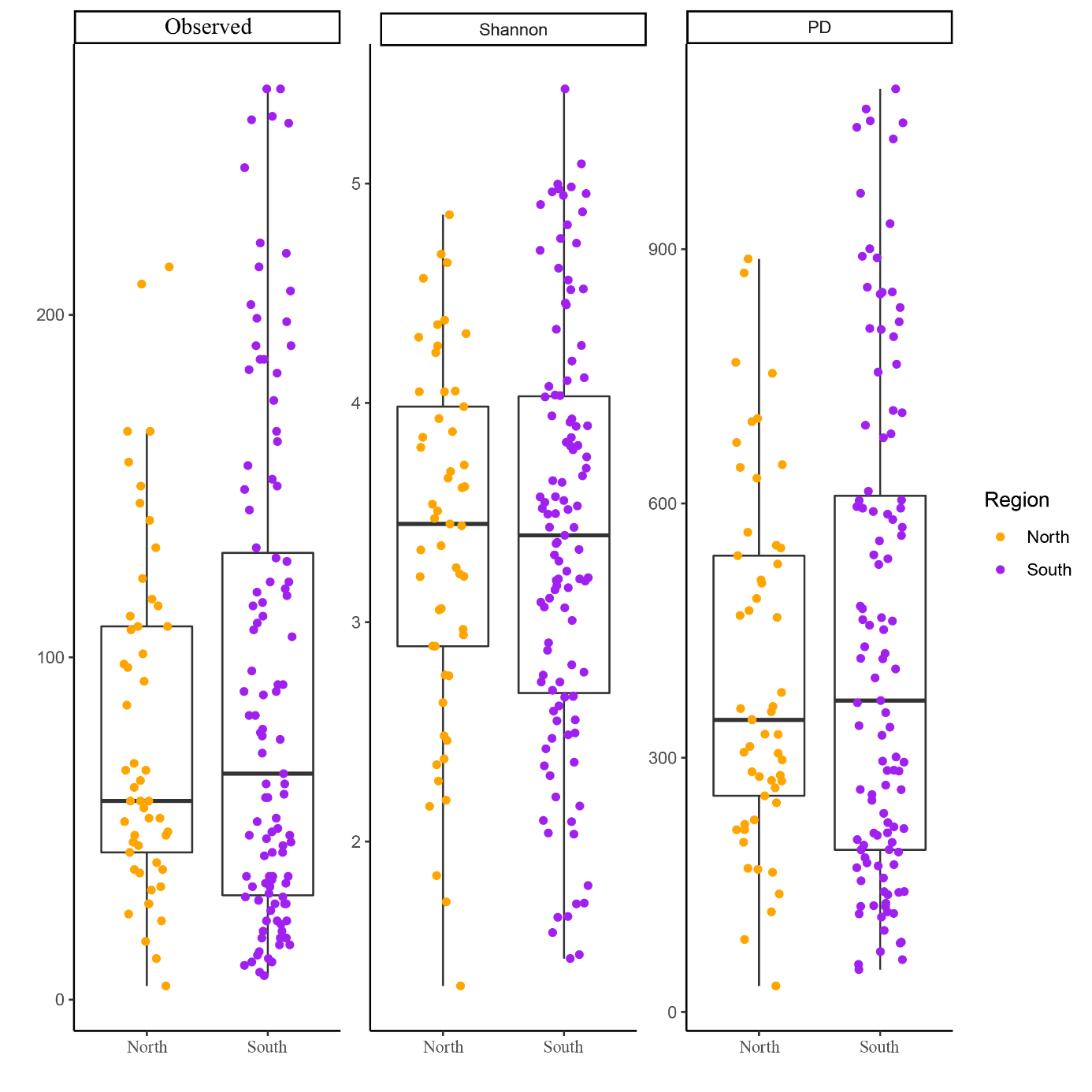

**Figure S6**. A non-metric multidimensional (NMDS) plot (stress value=0.13) of weighted and unweighted UniFrac distances in skin bacteria beta diversity matrices for moor frog (*R.arvalis*) between the northern (Luleå) and Southern (Uppland) region (PERMANOVA; Adonis2, p<005). In all charts, orange indicates samples from the north whereas purple indicates samples from the south.

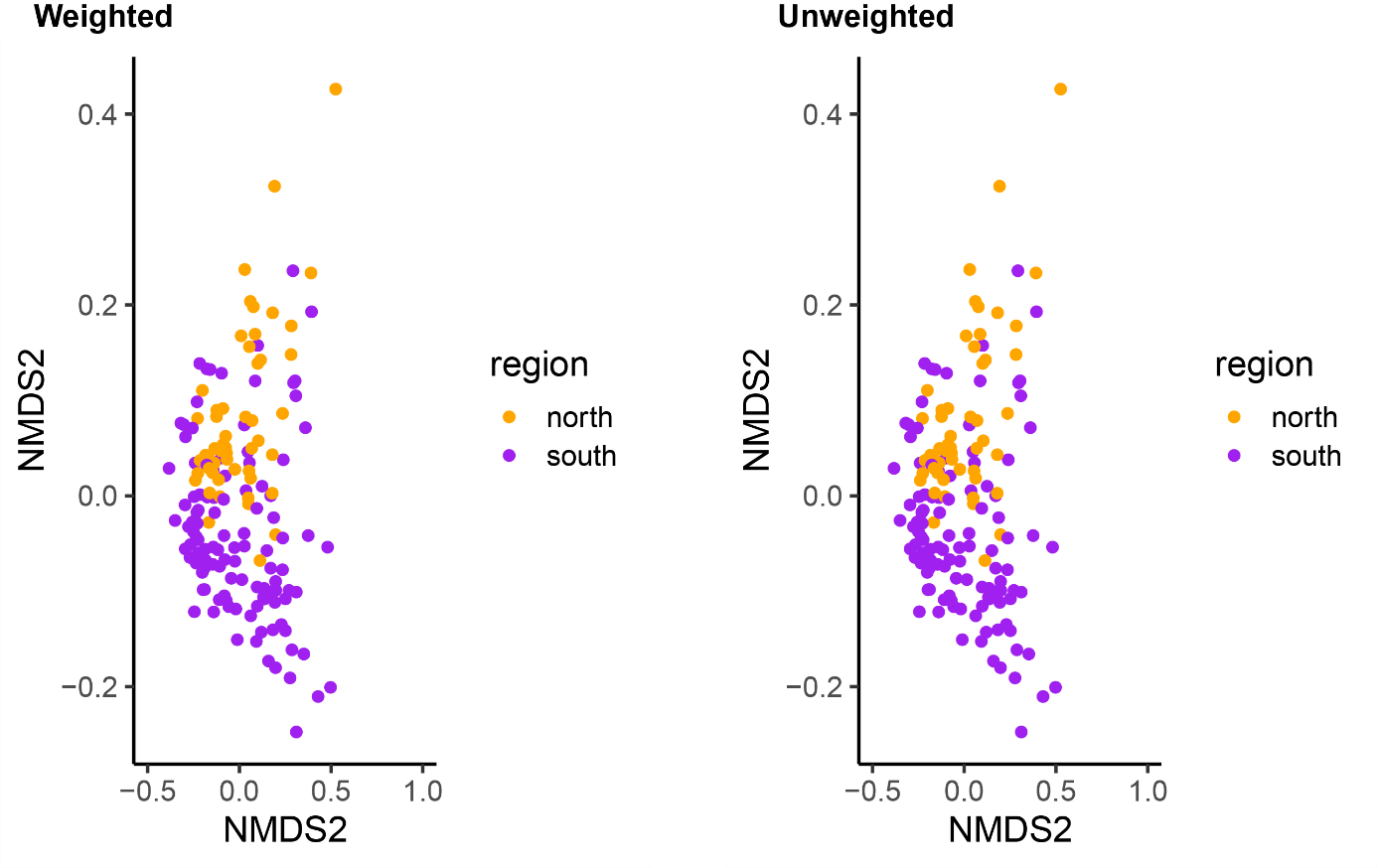

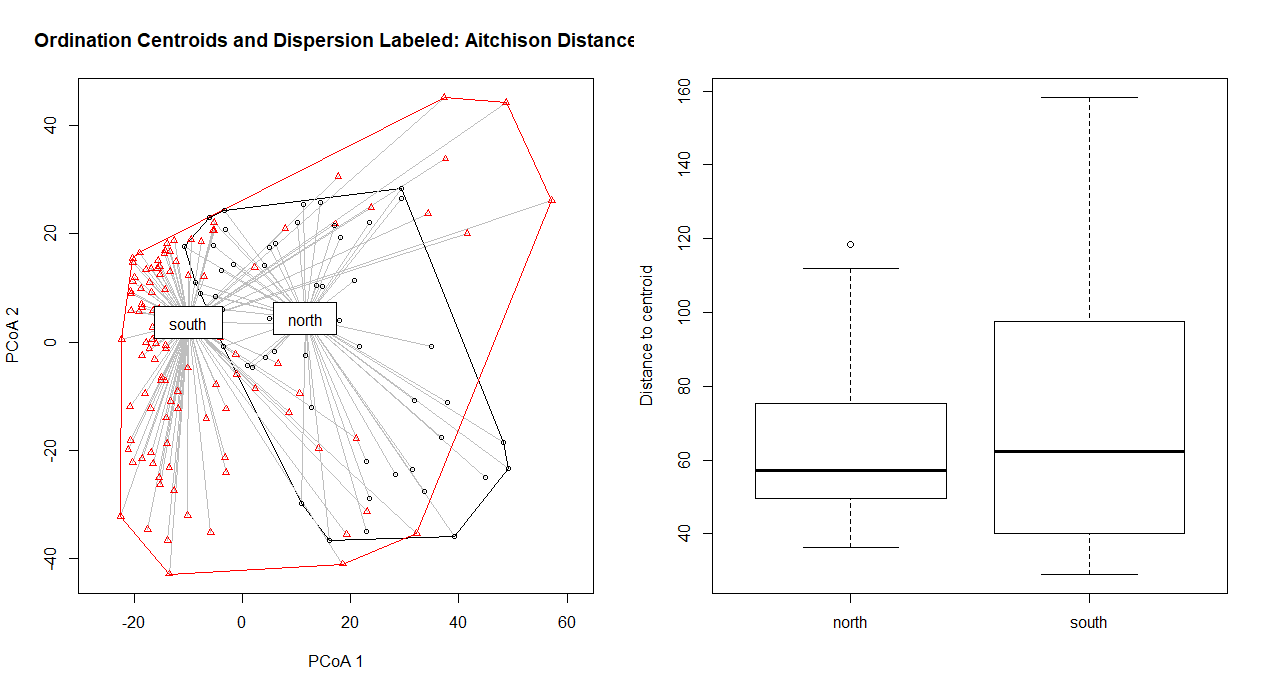
**Figure S7** A) PCoA plot demonstrating the beta diversity of different southern and northern cluster at ASVs level on weighted UniFrac distance derived from 16S microbial sequences (PERMANOVA; Adonis2, P <0.05). B) Represent the distance to the polygon centroids (PermDISP; p>0.05).

**B**

**A**

**Figure S8**. Shannon diversity index in relation to MHC heterozygosity for multiple regression on matrix (MRM) test. Each point represent a pairwise distance between the Shannon diversity index and the MHC heterozygosity. North-to-north comparisons are colored in orange, south-to-south comparisons are colored in purple and north-to-south comparisons are colored in grey.

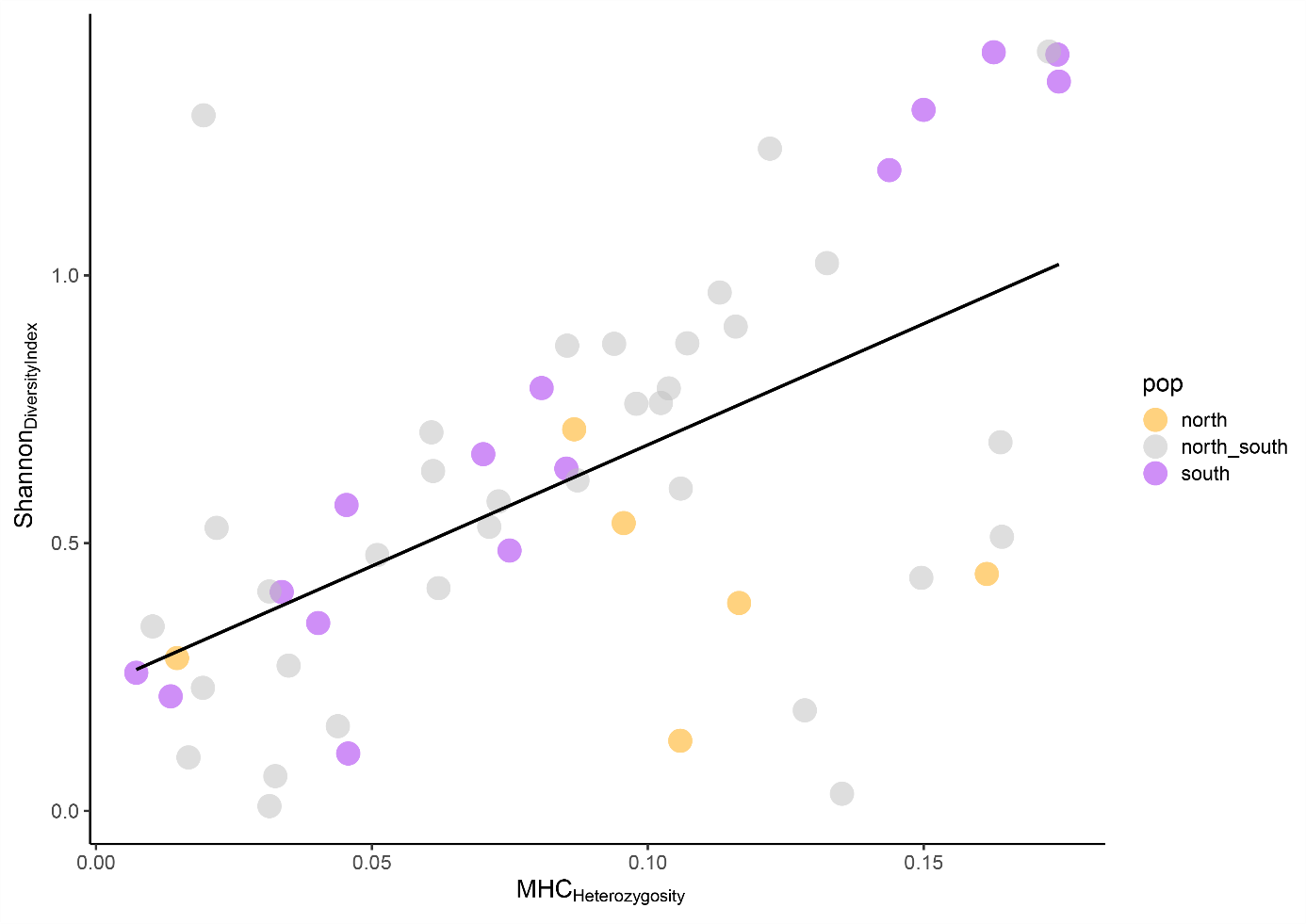

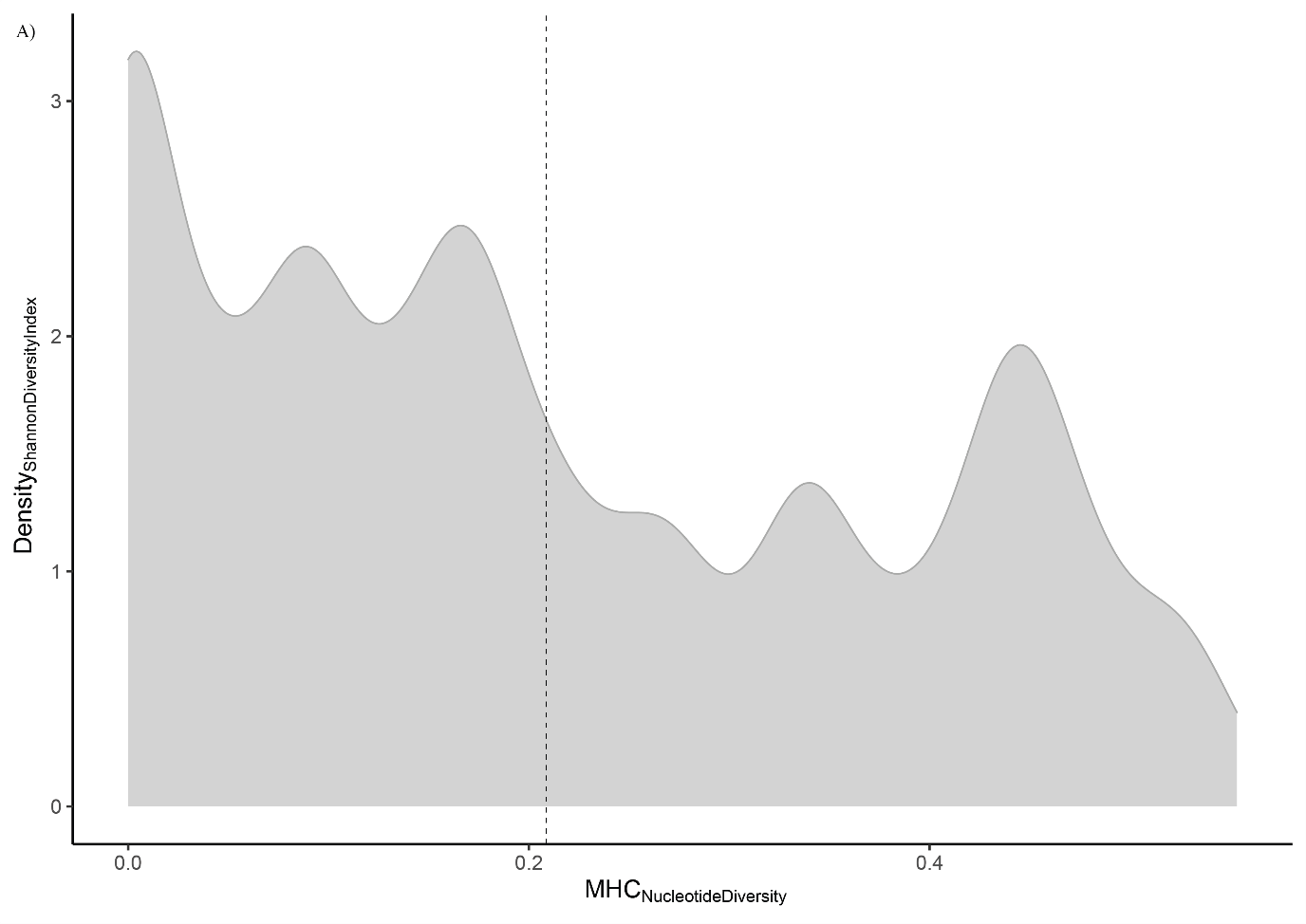
**Figure S9**. Density plot describing the Shannon diversity index in relation to the MHC nucleotide diversity. The black dashed line shows the overall trend in diversity.

**Figure S10.** Overall microbial diversity (Shannon diversity index) in individuals belonging to the group “DistinctS” and grouped by region of origin. Individuals from Uppland (Uppsala) are represented in purple and individuals from Luleå are represented in orange.

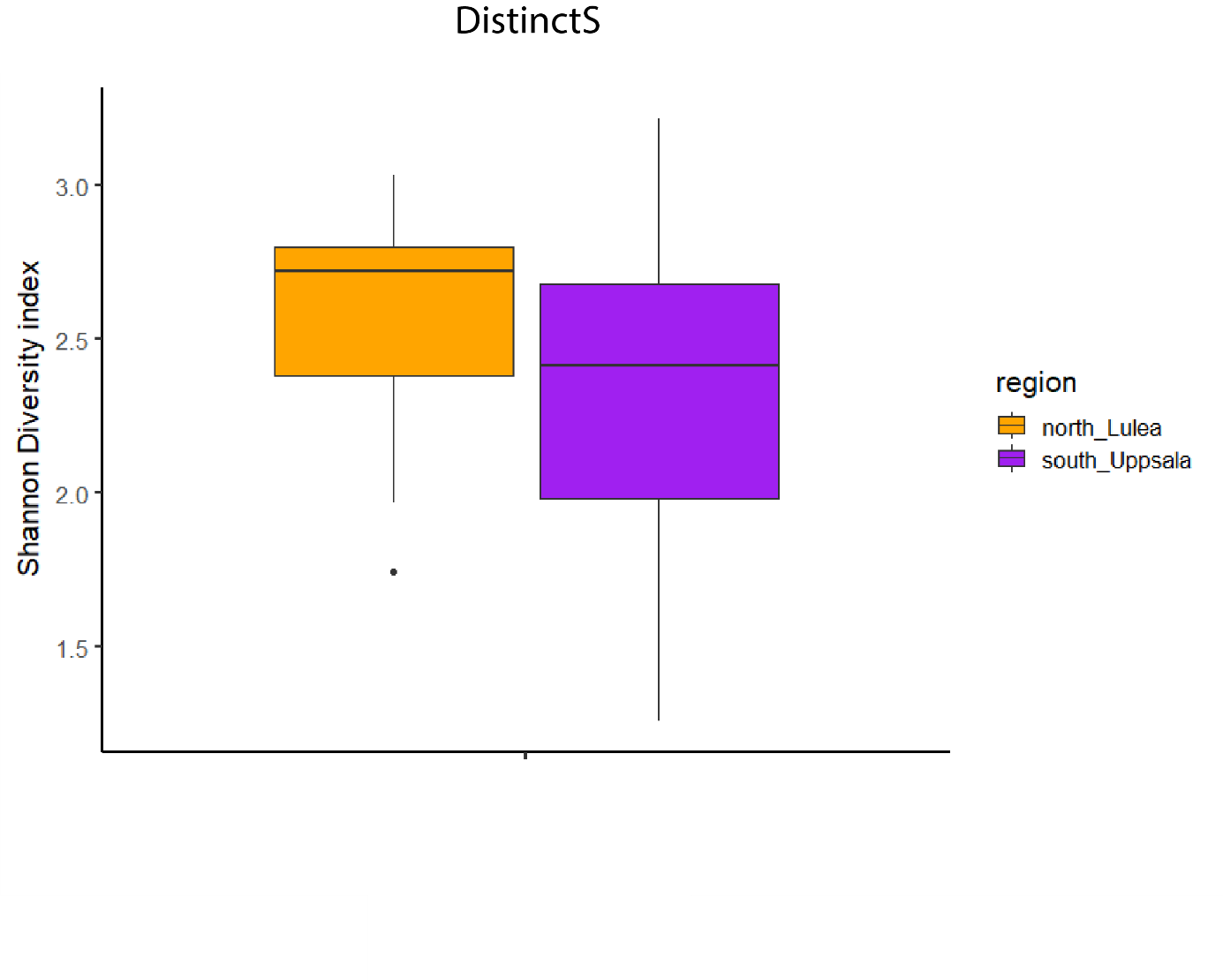

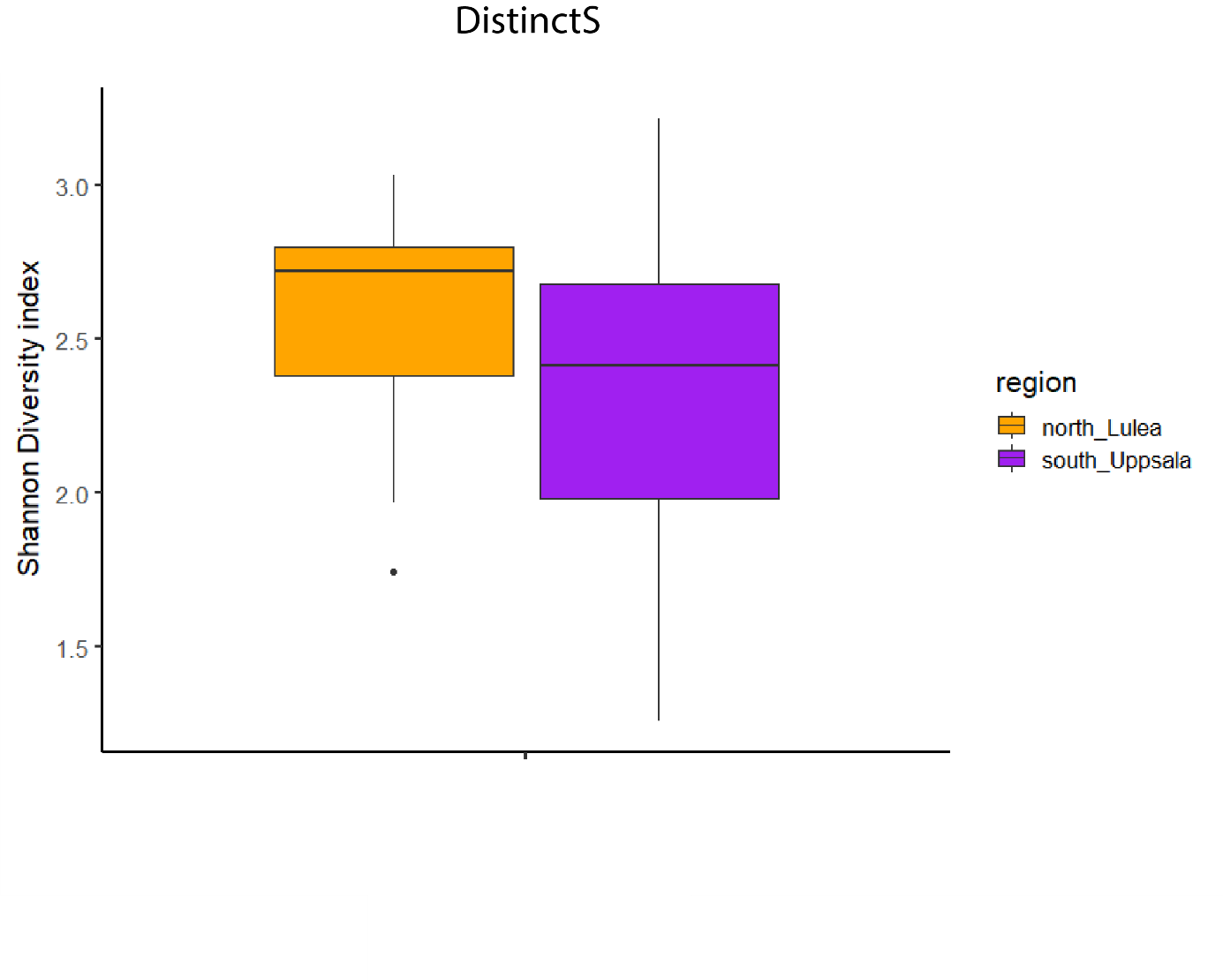

**Figure S11.** A) RDA plot showing the patterns of separations for all supertype genotypes characterized based on the MHC class II exon 2 data set. B) The RDA plots show only the significant supertype_genotypes that carry a specific skin bacterial composition based on the PERMANOVA analyses (Adonis; p<0.05), where S1_3 is colored in purple, S2_2 is colored in light pink, S3_3 is colored in red and S2_4 is colored in green.

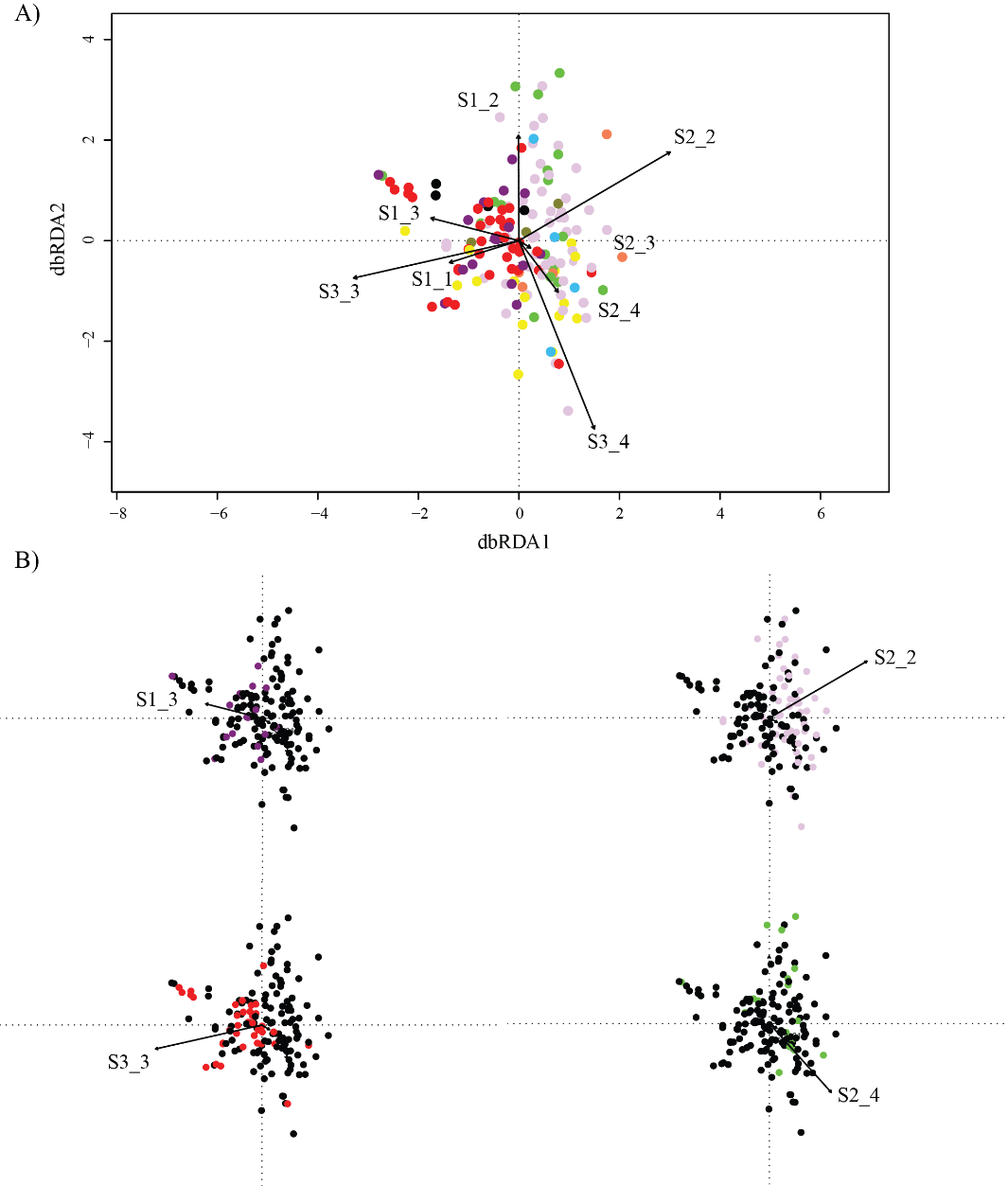

**Figure S12**. Taxa count base on the family classification per supertypes. With and (*) are represented the significant different ASVs based on the DEseq2 analyses that belong to a specific family between supertypes (S1, S2, S3 and S4) See Figure 7 for positive and negative correlations with MHC.

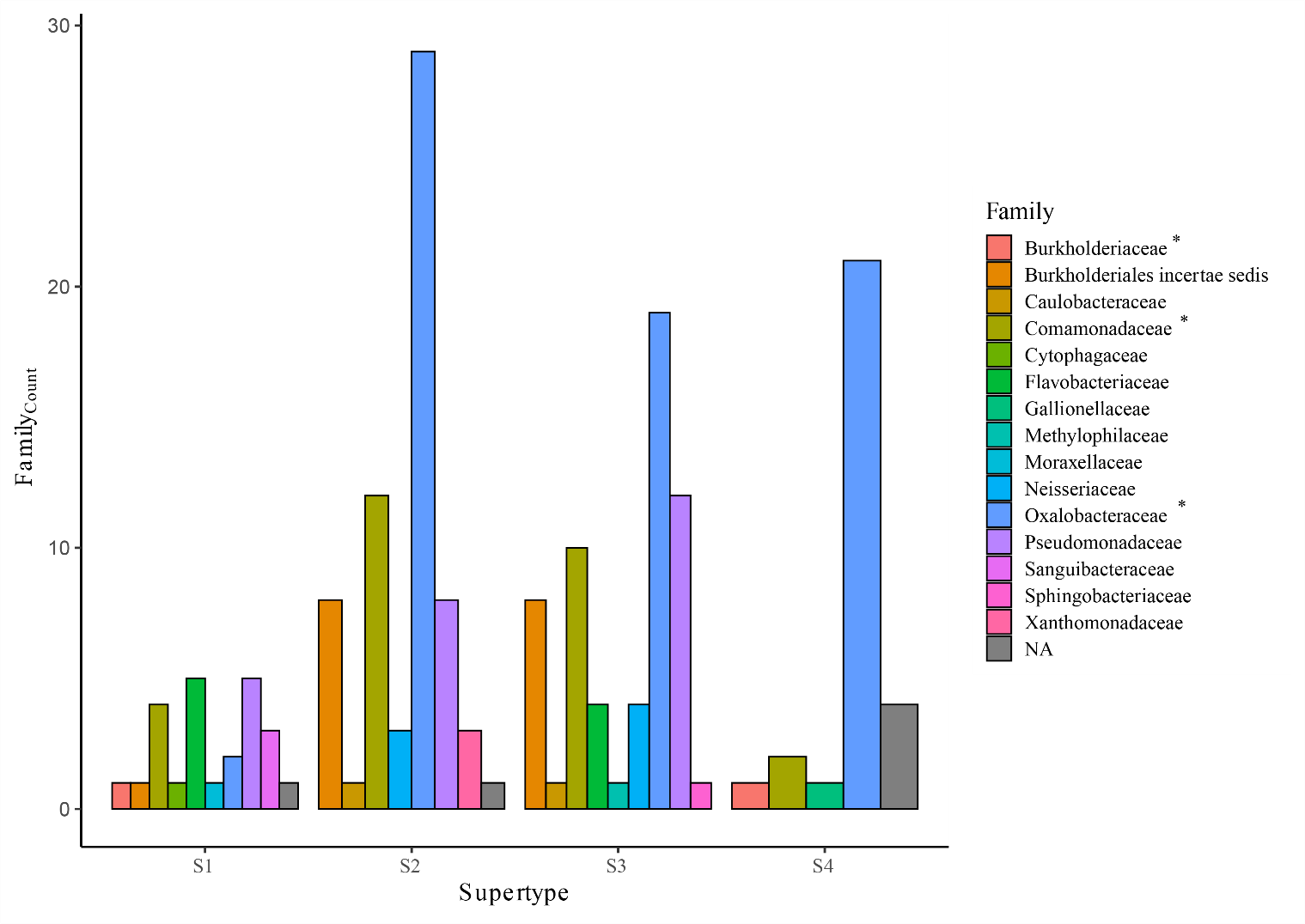
